# Pluripotent stem cell SOX9 and INS reporters facilitate differentiation into insulin-producing cells

**DOI:** 10.1101/2021.02.02.429390

**Authors:** Rabea Dettmer, Isabell Niwolik, Ilir Mehmeti, Anne Jörns, Ortwin Naujok

**Affiliations:** Institute of Clinical Biochemistry, Hannover Medical School, Hannover, Germany

## Abstract

Differentiation of human pluripotent stem cells into insulin-producing stem cell-derived beta cells harbors great potential for research and therapy of diabetes. The SOX9 gene plays a crucial role during development of the pancreas and particularly in the development of insulin-producing cells as SOX9+ cells form the source for NEUROG3+ endocrine progenitor cells. For the purpose of easy monitoring of differentiation efficiencies into pancreatic progenitors and insulin-producing cells, we generated new reporter lines by knocking in a P2A-H-2K^k^-F2A-GFP2 reporter genes into the *SOX9* locus and a P2A-mCherry reporter gene into the *INS* locus mediated by CRISPR/CAS9-technology. The knock-ins enable co-expression of the endogenous genes and reporter genes, report the endogenous gene expression and enable the purification of pancreatic progenitors and insulin-producing cells using FACS or MACS. Using these cell lines we established a new differentiation protocol geared towards SOX9+ cells to efficiently drive human pluripotent stem cells into glucose-responsive beta cells.

## Introduction

The SOX9 protein belongs to a large family of high-mobility domain transcription factors with pleiotropic functions during vertebrate development, cellular maintenance and disease development [1]. In humans SOX9 haploinsufficiency leads to campomelic dysplasia with pancreatic dysmorphogenesis [2]. This evidence and other results from several transgenic mouse models [3, 4] have led to the conclusion that SOX9 belongs to the group of master regulators of pancreatic development [1].

SOX9 expression during the initial forming of the dorsal and ventral pancreatic buds strongly co-localizes with PDX1 in mouse and man [4, 5]. In mice, when the early unpolarized epithelium branches into a plexus with proximal trunk and distal tip domains, Sox9 is expressed co-localized with Cpa1 and Pdx1 in the tip domain or with Nkx6.1 and Pdx1 in the trunk domain [4]. The distal tip domain is considered as the cellular pool for acinar differentiation, whereas the proximal trunk domain harbors the development niche for bipotent ductal/endocrine precursor cells [6]. Bipotent precursor then give rise to Neurog3+ cells. Later, during mid second transition, Sox9 expression is receding from the tip domain and becomes restricted to the proximal trunk cells. By late gestation and in adults Sox9 expression is confined to centroacinar and ductal cells [1, 4].

Thus, SOX9 is an interesting target for the generation of new human pluripotent stem cell (hPSC) reporter lines in order to target the development niche that gives rise to the endocrine lineage. This would open up new opportunities to develop efficient differentiation protocols for the generation of stem cell-derived beta cells or organoids, respectively (SC-derived beta cells/SC-derived organoids). Therefore, the aims of this study were to generate two types of reporter cell lines, namely a *SOX9* and a *SOX9*/*INS* knock-in cell line and a differentiation protocol optimized towards the generation of SOX9+ MPCs.

For that purpose, we used CRISPR/Cas9 to knock-in reporter genes, GFP2 and the surface antigen H-2K^k^, in frame by homology directed repair (HDR) into the *SOX9* locus and additionally mCherry into the *INS* locus. This permits monitoring and cell purification of *SOX9* and *INS* expressing populations.

We can show that PDX1+ pancreatic-duodenal cells described in earlier studies [7, 8], effectively differentiate into SOX9+ multipotent pancreatic progenitor cells (MPCs) with co-expression of CPA1 or NKX6.1. By use of this triple knock-in cell line we could track the conversion of SOX9 MPCs into SC-derived beta cells. In parallel, we have developed a 3D differentiation protocol to efficiently generate a large number of SC-derived organoids predominantly composed of monohormonal beta cells.

## Results

In order to generate reporter PSC lines, a knock-in strategy based on CRISPR/Cas9-induced DSBs was developed (Supplementary Table 1). A repair vector comprising 500 bp 5’ and 3’ homology arms, two reporter genes, namely H-2K^k^ and GFP2, separated by 2A cleavage sites, and a floxed selection gene cassette was cloned (Fig 1A). After nucleofection and selection by antibiotics, pluripotent stem cell colonies with typical morphology were expanded and genotyped by PCR. Correct insertion was verified by sequencing. The HES-3 clone SC30 and the Phoenix iPSC clone NSC20, both with homozygous integration, were selected for further work. Our optimized endoderm differentiation protocol [9] robustly produced > 90% CXCR4-positive cells, which were also predominantly positive for the anterior foregut endoderm markers CD177 and CD275 as recently described [10] (Supplementary Fig. 1A-D). Then six different protocols for differentiation of endoderm into MPC were tested (Supplementary Fig. 2). Protocol # 6, which is based on our protocol for the differentiation of hPSC into PDX1+ pancreatic-duodenal cells (Supplementary Fig. 1E) [7, 8] expanded by Stage 3 and Stage 4 media from [11] was then selected and termed 2D experimental differentiation protocol (Fig. 1B). This 4-stage protocol, when applied to SC30 and NSC20 cell clones, yielded in an expression peak of ∼40-50% GFP2+ cells between day 10 and 13 of differentiation (Fig 1C). GFP2+ positive cells additionally expressed H-2K^k^, thereby allowing cell purification by either FACS or MACS (Fig 1D/Supplementary Fig. 3). Next, GFP2+/GFP2-cells were sorted by FACS and analyzed upon expression of MPC marker genes. GFP2+ cells expressed significantly more *HNF6, NKX6-1, PDX1* and *SOX9* compared to GFP2-cells. Only *NEUROG3* was stronger expressed in GFP-cells (Fig. 1E/Supplementary Fig. 4).

**Figure 1.**
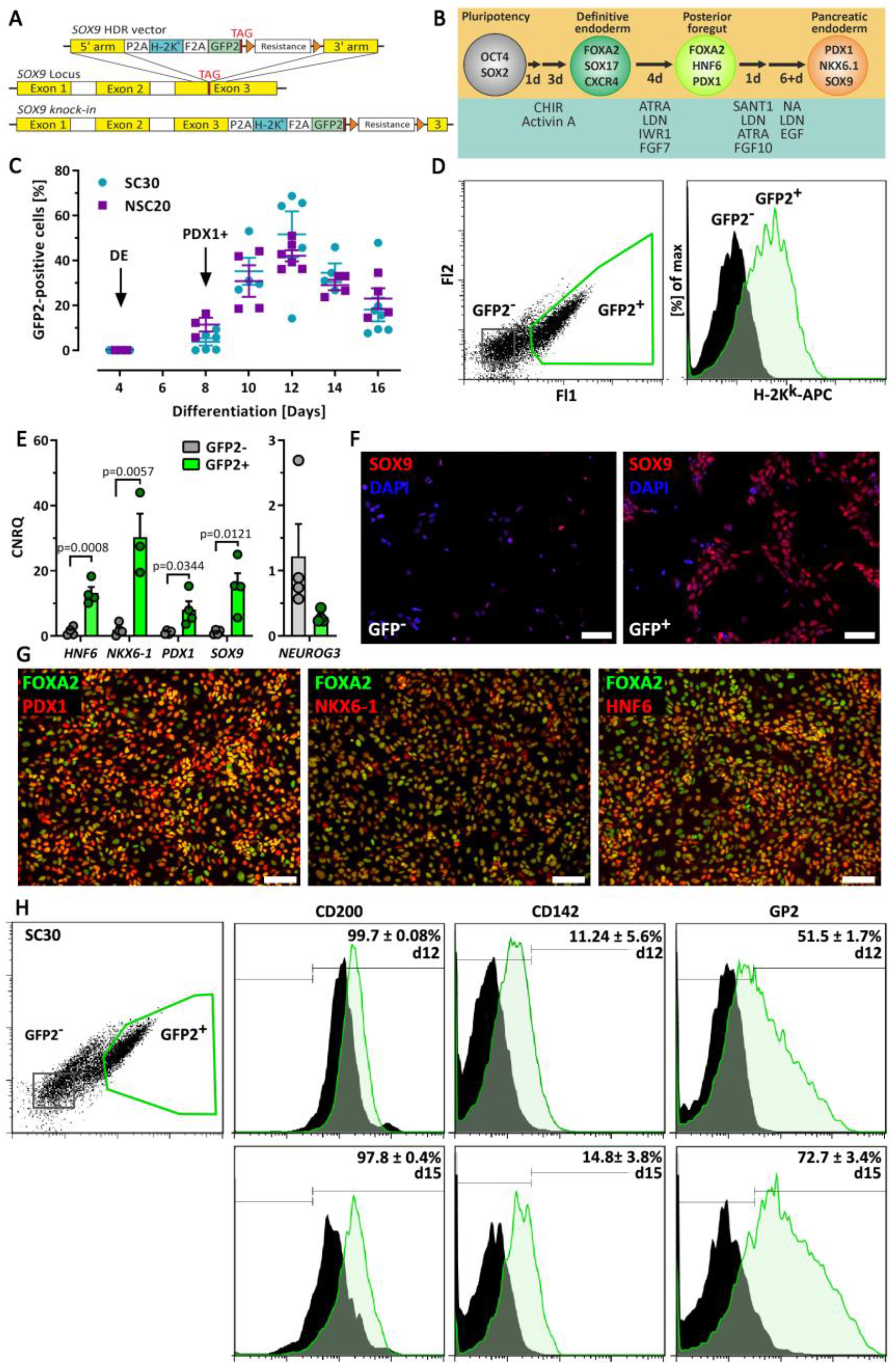
(**A**) Schematic presentation of the SOX9 HDR vector and the human *SOX9* locus before and after homologous recombination. (**B**) Schematic presentation of the 4-stage experimental 2D differentiation protocol for the generation of SOX9+ progenitors. (**C**) GFP2 expression during differentiation of the SC30 and NSC20 cells. Data are means ± SEM, n= 4-8. Arrows mark developmental stages. (**D**) Flow cytometry dot plot and histogram of SC30 GFP2 expression and gated H2-K^k^ staining at d12. (**E**) RT-qPCR analysis of sorted SC30 derived GFP2+ and GFP2-cells (d11-d12). Depicted is the relative gene expression of *HNF6, NKX6-1, PDX1*, and *SOX9*. Data are means ± SEM, n= 3-4. Two-tailed *Student’s* t-test. (**F**) Immunofluorescence staining of SOX9 (red) in SC30 derived GFP2^+^ and GFP2^-^ cells (d12). Nuclei were counterstained with DAPI. (**G**) Immunofluorescence staining of PDX1, HNF-6 and NKX6-1 (red) and FOXA2 (Green) of sorted SC30 derived GFP2^+^ cells (d12). (**F**/**G**) Scale bar = 100 µm. (**H**) Flow cytometry dot plot and histogram of SC30 GFP2 and CD200, CD142 and GP2 expression at d12 and d15 (end of stage 4). Bifurcated gates were set according to unstained controls, values are means ± SEM, n = 3-4.

SOX9 protein predominately occurred in the nucleus of GFP2+ cells and was low or absent in GFP2-cells (Fig 1F). Consistently to the gene expression data, PDX1, NKX6-1 and HNF6 were prominently expressed in GFP2+ cells (Fig 1G). Next, surface markers described to be expressed on MPCs, namely CD200 and CD142 [12] and GP2 [13], were measured by flow cytometry. D12 and d15 GFP2+ cells expressed CD200 and GP2 and faintly CD142, but only GP2 was differently expressed compared to GFP-cells (Fig. 1H).

According to the spatiotemporal expression pattern of Sox9 during pancreas organogenesis in mice, we expected to detect CPA1+/SOX9+ and NKX6.1+/SOX9+ cells as representatives of distal and proximal tip/trunk cells in this differentiation model. Western blot analysis showed a peak CPA1 expression in d15/d18 cells. To confirm this finding we used MACS to purify SOX9 MPC and further differentiated them in 2D using the stage 5-7 media described by Pagliuca and co-workers [14]. Re-seeded and purified SOX9 MPC formed tight clusters/colonies after cell sorting. At d15 mosaic NKX6.1 and CPA1 expressing cells were readily detected and by d18 NKX6.1 was almost homogenously expressed in these colonies. D18 marked also the expression peak of scattered NEUROG3+ and very few CPA1+ cells (Fig. 2C). The detection of the first NEUROG3+ cells coincided with insulin and C-peptide+ cells, which from d18 grew out in clusters. These structures were embedded in CK19+ epithelial colonies. Gene expression changes of sorted and further differentiated cells are presented in Supplementary figure 5. We next analyzed insulin and C-peptide content and whether these SC-derived beta cells would release insulin. We also compared sorted vs non-sorted differentiation experiments. Cell sorting yielded in a ∼4-fold increase of insulin and C-peptide content compared to non-sorted cells.

**Figure 2.**
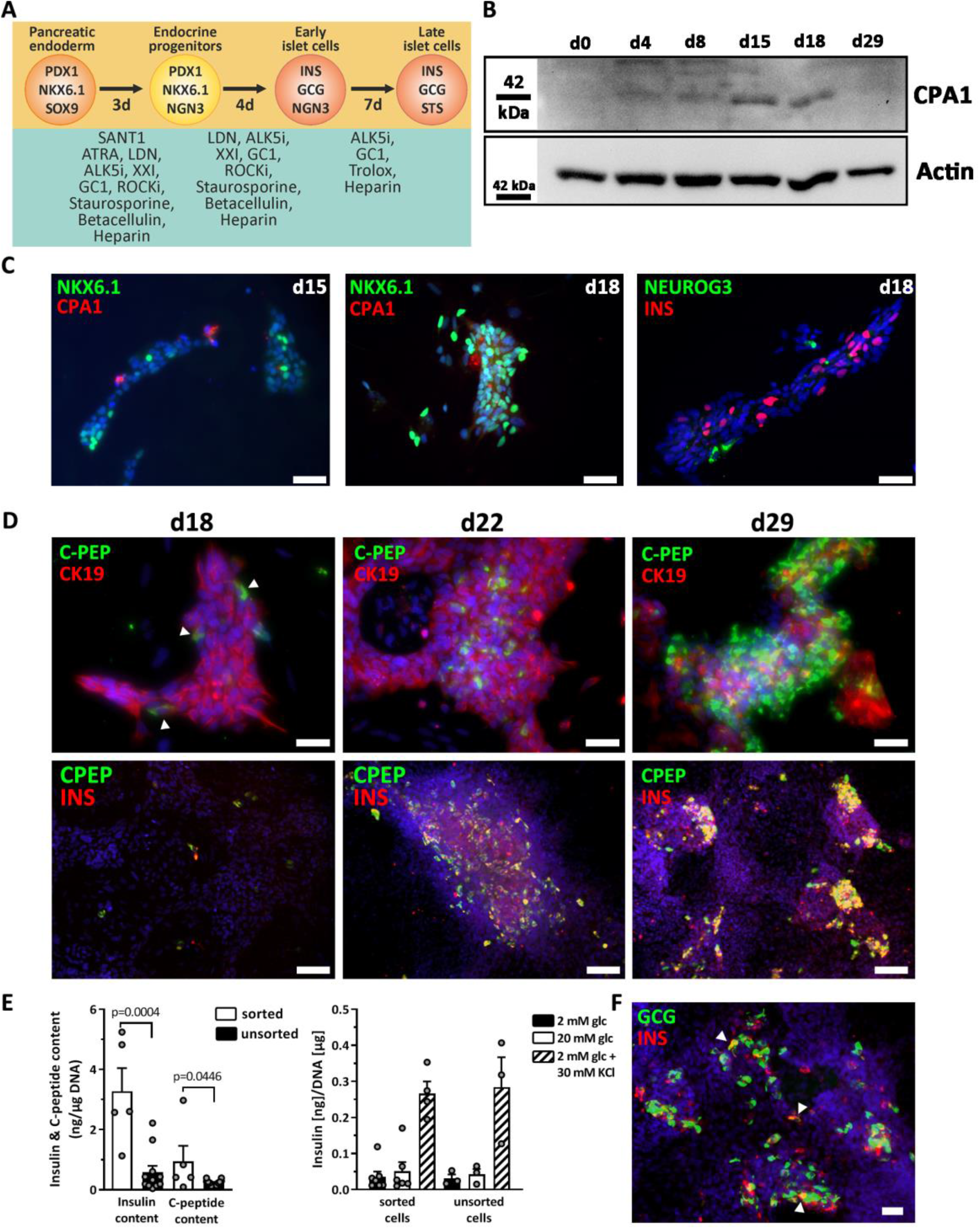
(**A**) Scheme of the 3-stage protocol for differentiation of SOX9+ MPCs into stem cell-derived beta cells. MACS was performed at d12 using the SC30 cell clone. (**B**) Analysis of CPA1 protein expression during differentiation (d0-d29) by Western Blot. (**C**) Immunofluorescence staining of NKX6.1/CPA1 and NEUROG3/insulin after stage 4 (d15) or 5 (d18), respectively. Scale bar = 50 µm. (**D**) Immunofluorescence staining of CK19/C-peptide and C-peptide at d18, d22 or d29 of differentiation. Scale bar = 50 µm (CK19/C-peptide) or 100 µm. Arrowheads indicate early insulin-positive cells. (**E**) Measurement of insulin and C-peptide secretion and content in d29 sorted SC30 cells vs unsorted cells. Data are means ± SEM, n= 5-12 (content) and n=3-7 (secretion). Two-tailed *Student’s* t-test, *** p < 0.001, * p < 0.05. (**F**) Immunofluorescence staining of glucagon and insulin in d29 cells. Arrowheads mark polyhormonal cells. Scale bar = 50 µm.

However, insulin release was only triggered by KCl and not by elevated glucose concentrations and mean variation was high (Fig 2E). Since differentiation of hPSCs into polyhormonal SC-beta cells is a common phenomenon, we double stained insulin and glucagon and found to a minor extent polyhormonal cells with both hormones (Fig. 2F).

Next, to take advantage of the SOX9 reporter cells, we tested the impact of stage 3 and stage 4 media components. In a subtraction assay we compared GFP2 expression in control cells cultured for 24 h in stage 3 and 72 h in stage 4 against treatments, where stage 3 was omitted or individual components of stage 4 were left out (Fig 3A). The omission of stage 3 showed the greatest and significant reduction of GFP2+ cells. In descending order, the number of GFP2+ cells was also reduced by omitting LDN, EGF and NA from the stage 4 medium. Addition of a PKC activator as used in some differentiation protocols, however, did not increase the number of GFP2+ cells in our protocol. Then we titrated the amount of EGF in stage 4 medium and were able to determine that the maximum of GFP2 expression was reached for both cell lines from ∼100 ng/ml EGF and on.

**Figure 3.**
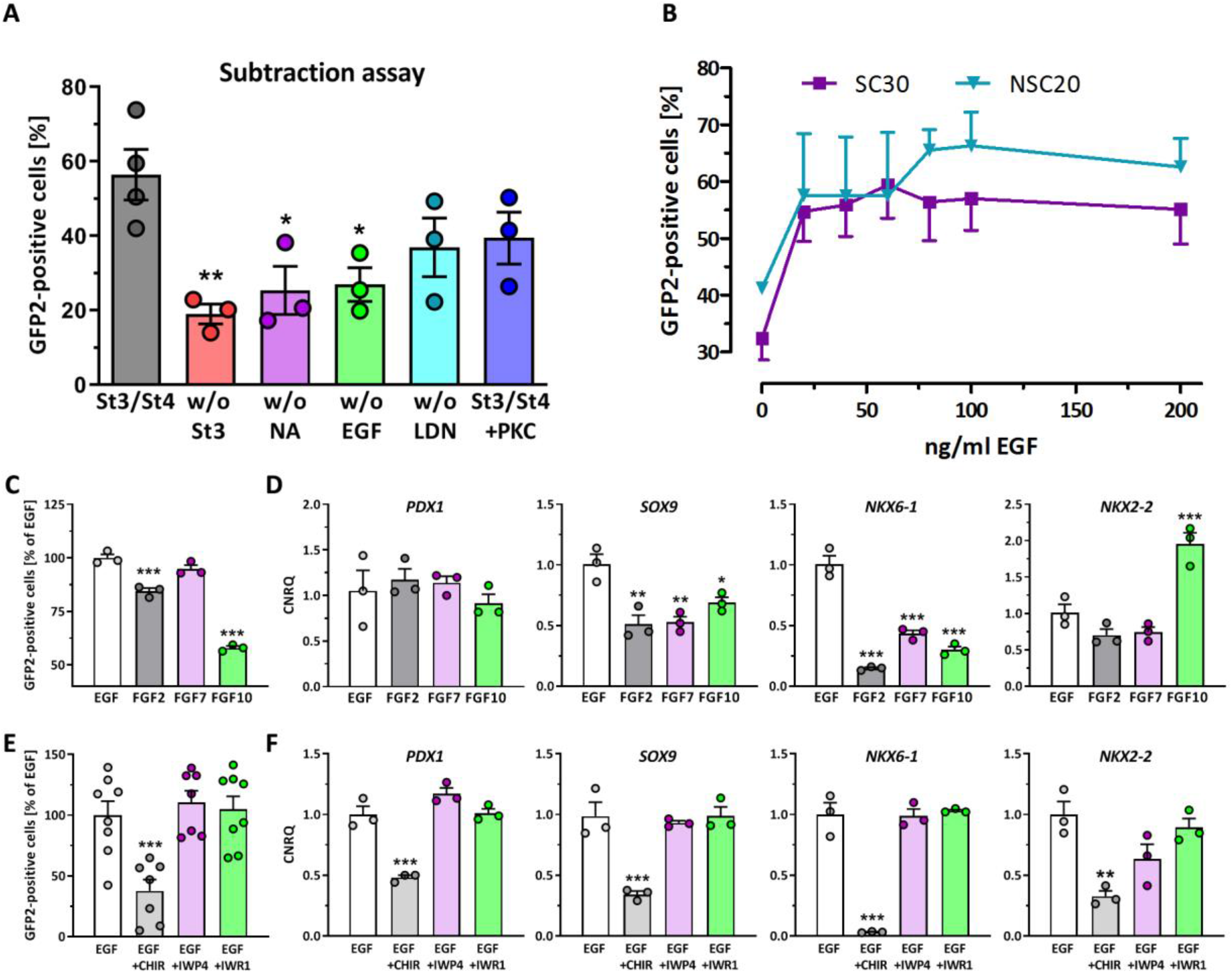
(**A**) Measurement of GFP2+ cells at d12 after subtraction of either stage 3, nicotinamide (NA), EGF, or LDN193189 (LDN). For comparison stage 4 plus protein kinase C activation by 100 nM PDBu. Data are means ± SEM, n= 3. (**B**) GFP2-expression in dependence of the EGF concentration at d12. Data are means ± SEM, n= 3. (**C/D**) Effect of different growth factors each used at 100 ng/ml on GFP2 expression (**C**) at d12 and pancreatic marker gene expression (**D**). Depicted is the relative gene expression of *PDX1, SOX9, NKX6-1*, and *NKX2-2*. Data are means ± SEM. n= 7-8 (GFP2 flow cytometry), n=3 (RT-qPCR). (**E**/**F**) Effect of canonical Wnt-signaling on GFP2 expression (**E**) at d12 and pancreatic marker gene expression (**F**). The pathway was activated by CHIR (3 µM) or inhibited by IWP4 (1 µM) or IWR-1 (2 µM). Depicted is the relative gene expression of *PDX1, SOX9, NKX6-1*, and *NKX2-2*. Data are means ± SEM. n= 3 (GFP2, RT-qPCR). ANOVA plus *Dunnett’s* post-test, *** p < 0.001, ** p < 0.01, * p < 0.05.

The SOX9 gene is controlled, amongst other mechanisms, by the FGFR2b signaling pathway and by Wnt/beta-catenin signaling [15]. Therefore we compared the effects of the growth factors FGF7 and FGF10, which signals through FGFR2b, and of EGF and FGF2 used in other beta cell differentiation protocols at this stage of differentation [11, 13, 14, 16]. Also the effect of Wnt/beta-catenin inhibition and activation was analyzed (Fig 3C-F). Using the SC30 cell clone, it turned out that the incubation with 100 ng/ml EGF produced the highest number of GFP2+ cells and showed the strongest expression of *PDX1, SOX9* and *NKX6-1*. FGF10 and FGF2 at the same concentration caused a significant decrease of GFP2+ cells compared to EGF (Fig. 3C). Wnt/beta-catenin activation of this pathway by CHIR99021 showed a significant inhibitory effect both on the number of GFP2+ cells and on the expression of *PDX1, SOX9, NKX6-1* and *NKX2-2*. Addition of the inhibitors of Wnt/beta-catenin signaling IWP4 and IWR-1 did, however, not yield in a further increase of GFP2+ cells or enhanced gene expression (Fig 3E/F). The same pattern was found for the NSC20 cell clone (Supplementary Fig. 6A/B). Furthermore, we were able to determine a significant increase in GFP2 expression from 42% to 74% for the SC30 cell clone and from 46% to 77% for the NSC20 cell clone after upscaling from 2D differentiation in 12-well cavity format to 10 cm culture dishes (Supplementary Fig. 6C).

For simplification of the monitoring process, we knocked-in mCherry into the human *INS*-locus of the SC30 cell clone. This allowed us to measure SOX9 progenitor generation and their conversion into SC-derived beta cells using flow cytometry. For this, a repair vector similar to the repair vector for the *SOX9*-locus was cloned. It comprised 500 bp 5’ and 3’ homology arms, the reporter gene mCherry and a floxed selection gene cassette (Fig 4A). For the knock-in into the *INS* locus, CRISPR/Cas9 nickases were used (Supplementary Table 1). After verification of correct insertion by sequencing, the homozygous SC30 ICNC4 cell clone was selected for further work.

**Figure 4.**
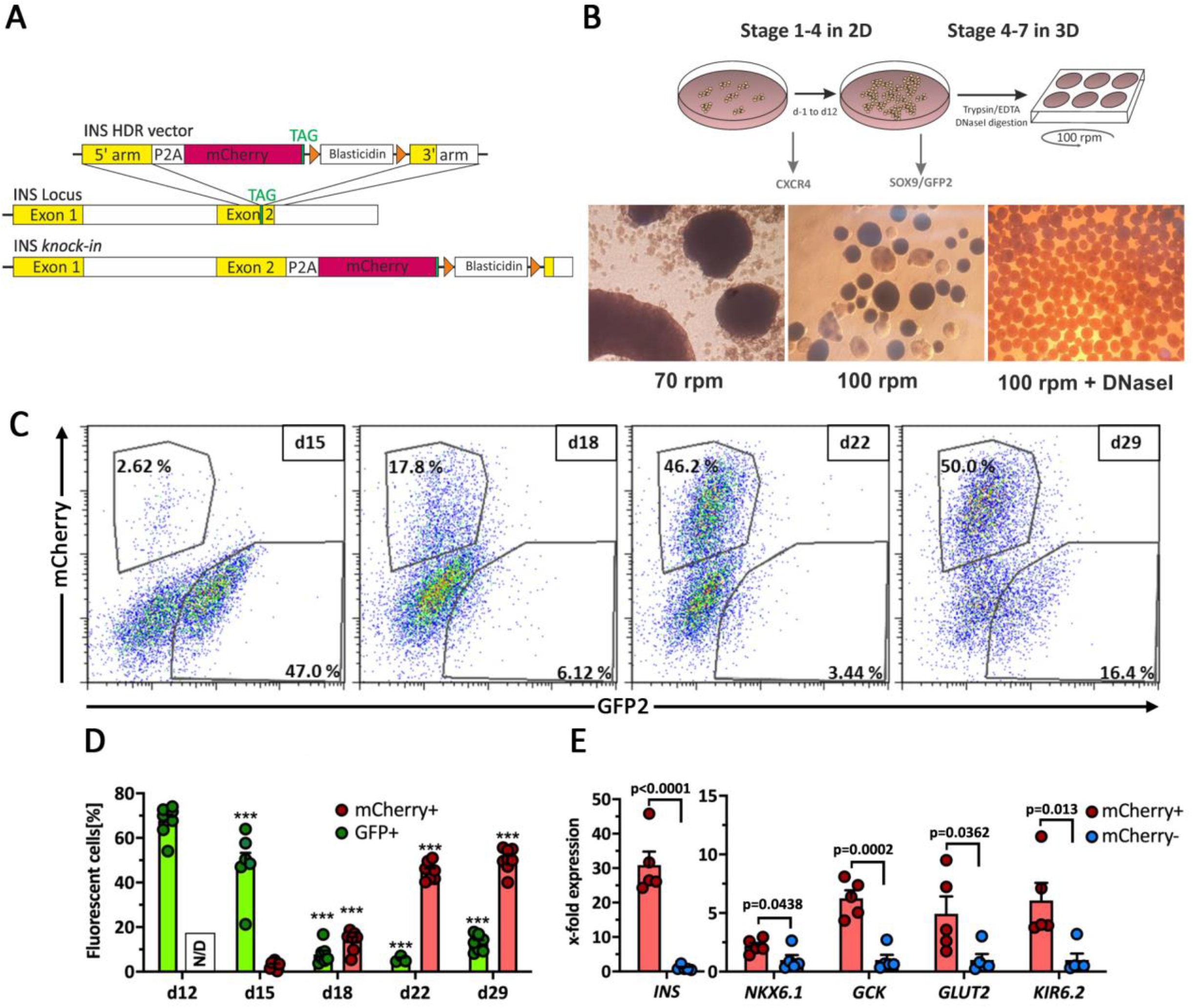
(**A**) Schematic presentation of the INS HDR vector and the human *INS* locus before and after homologous recombination. (**B**) Schematic presentation of the 3D production protocol for the differentiation of SC30/NSC20 and SC30 ICNC4 cell clones into stem cell-derived islets and phase contrast images of clusters generated by 3D orbital shaking culture. Monitoring of differentiation with CXCR4 and GFP2 measurement on d4 and d12. (**C**) Representative flow cytometry dot plots of mCherry vs GFP2 protein expression during 3D differentiation. The numbers in the gates indicate the percentage of SOX9- or INS-expressing cells at different stages of differentiation. (**D**) Kinetics of GFP2 and mCherry protein expression in SC30 ICNC4 cells during 3D differentiation. Values are means ± SEM. n= 3-7. ANOVA plus Tukey’s post-test, *** p < 0.001, ** p < 0.01, * p < 0.05, compared to d12/d15 of differentiation. (**E**) RT-qPCR analysis of the beta cell marker genes *INS, NKX6-1*, glucokinase (*GCK*), *GLUT2* and *KIR6*.*2* in sorted mCherry+ vs mCherry-cells at d29 of differentiation generated with the SC30 ICNC4 cell clone. Values are means ± SEM. n= 5, *Student’s* t-test *** p < 0.001, * p < 0.05. Expression values for mCherry-cells were set to 1.

With recent reports on the improvement of differentiation and maturity through 3D culture we adapted our 2D experimental protocol and introduced 3D orbital shaking from d12 on until the end of differentiation (Stage 4-7). Stage 3 and 4 media were adapted according to our previous findings. For 3D differentiation, d12 cells were gently dissociated and transferred to 6-well suspension culture dishes on an orbital shaker. Different rotating speeds from 70 rpm to 100 rpm were tested. Finally, we settled with 100 rpm and around 2 x 10^6^ cells per ml (5 ml per cavity). To reduce cell clumping the cells were treated with DNase I before seeding, yielding in equally formed, round and small spheroids 24 h after transition from 2D to 3D culture. The spheroids were kept in culture for an additional 16-17 days without resizing. (Fig 4B, Supplementary Fig. 7).

During this time the kinetics of GFP2 and mCherry expression (*SOX9* and *INS* expression, respectively) were monitored by flow cytometry. It could be shown that the increase in *INS*+ cells was accompanied by a parallel decrease in *SOX9+* cells starting from d15 (Fig 4C/D). Next, we verified the *INS* knock-in by comparing gene expression of beta cell genes in mCherry+ vs mCherry-cells. *INS* gene expression was 31-fold higher in mCherry+ cells and other beta cell markers were between 2.2- and 6.1-fold higher expressed compared to mCherry-cells verifying functionality of the knock-in (Fig 4E). The gene expression kinetics of various pancreatic and endocrine genes was monitored during the 2D/3D differentiation process (Fig 5A). Of note here was again peak expression of *SOX9* and *CPA1* at d15 followed by peak *NEUROG3* expression three days later. In parallel with the peak in *NEUROG3* expression, the increase in islet cell hormones and *NKX6-1* could be recorded. Typical beta cell genes showed the same pattern in gene expression as insulin (Fig 5A).

**Figure 5.**
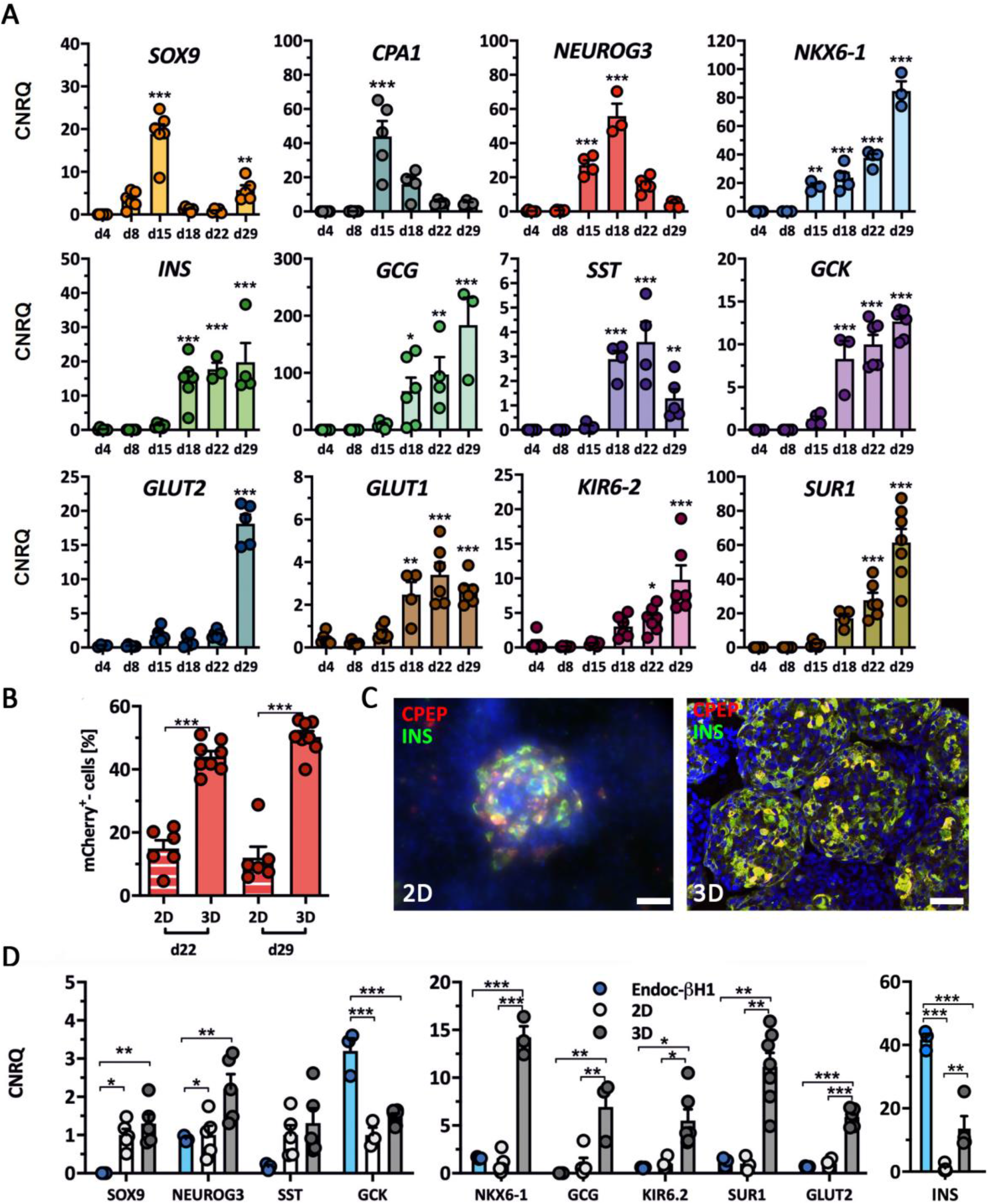
(**A**) Expression kinetics of pancreatic and endocrine genes during differentiation from d4 to d29 measured by RT-qPCR in SC30 ICNC4 cells. Values are means ± SEM. n= 3-6. ANOVA plus *Tukey’s* post-test, *** p < 0.001, ** p < 0.01, * p < 0.05, compared to d4 of differentiation. (**B**) Effect of 2D vs 3D differentiation on mCherry expression at d22 and d29 in SC30 ICNC4 cells. Values are means ± SEM. n= 4-6. ANOVA plus *Tukey’s* post-test, *** p < 0.001, ** p < 0.01, * p < 0.05. (**C**) Double-immunofluorescence staining of insulin (green) and C-peptide (red) in SC30-derived cells at d29 in 2D- or 3D-derived cells. Scale bar = 50 µm. (**D**) Effect of 2D vs 3D differentiation on pancreatic and endocrine marker gene expression measured by RT-qPCR comparing SC30 ICNC4 to EndoC-βH1 cells. Values are means ± SEM. n =3-6. ANOVA plus *Tukey’s* post-test, *** p < 0.001, ** p < 0.01, * p < 0.05. Relative expression values for 2D differentiation were set to 1.

Next, the number of *INS+* cells obtained in 2D culture was compared to 3D culture. 3D yielded in significantly more mCherry+ cells compared to 2D differentiation (mean 44% for d22 in 3D, 15% for d22 in 2D; mean 50% for d29 in 3D, 12% for d29 in 2D) (Fig 5B). The different efficiencies of 2D and 3D differentiation could also be observed after immunofluorescence staining of insulin and C-peptide (Fig. 5C). The expression of glucose recognition marker genes *GCK, KIR6*.*2, SUR1* and *GLUT2* and the transcription factor *NKX6-1* were significantly higher in 3D compared to 2D conditions and, except for *GCK* expression, comparable to EndoC-βH1 cells (Fig. 5D). Islet hormone expression was also higher in 3D conditions compared to 2D. Since EndoC-βH1 cells are a model cell line for human beta cells, the *INS* expression is higher compared to the heterogeneous composition of SC-derived organoids. The expression of the transcription factors *NEUROG3* and *SOX9* were also highest for 3D culture. This together with the gene expression analysis in 2D and 3D compared to EndoC-βH1 cells revealed the improvement of differentiation towards SC-derived organoids in 3D culture (Fig. 5D/Supplementary Fig. 8). The immunostaining of endocrine marker proteins, especially from beta cells, were then compared to those of d15 SC-derived spheroids and beta cells of islets and the surrounding tissue from human non-diabetic pancreas (Fig. 6). D29 SC-derived organoids generated from the SC30 cell clone were typically 200-300 µm in diameter and displayed a cytoplasmic co-localization of insulin and C-peptide in the majority of cells resembling the staining of beta cells in human islets. A polyhormonal staining of insulin or C-peptide with other islet peptides was rarely detected in these cells (Fig. 6/Supplemental Figure 9). Also the number of glucagon-positive cells was lower compared to 2D culture. NKX6.1 and PDX1 were in parallel localized in the nucleus of the insulin-positive cells in d29 SC-derived organoids. In comparison with human tissue, the only difference of the SC-derived organoids to human islets was the presence of some SOX9+ cells, whereas in human pancreas sections a clearer distinction between the SOX9-endocrine islets and SOX9+ exocrine parenchyma compartment was observed (Fig. 6). For comparison, immunofluorescence (IF) staining of insulin, C-peptide and glucagon of SC30 and NSC20 cells in 2D culture is depicted in Supplementary Fig. 10. Next the SC-derived organoids were further characterized. First we were able to measure a sharp increase in insulin and C-peptide content from d22 to d29 organoids (Fig. 7A). This content was comparable with EndoC-βH1 cells (101 ng insulin/µg DNA) and significantly higher comparing 2D with 3D (5.7 vs 302.27 ng insulin/µg DNA for SC30, respectively) (Fig. 7B).

**Figure 6.**
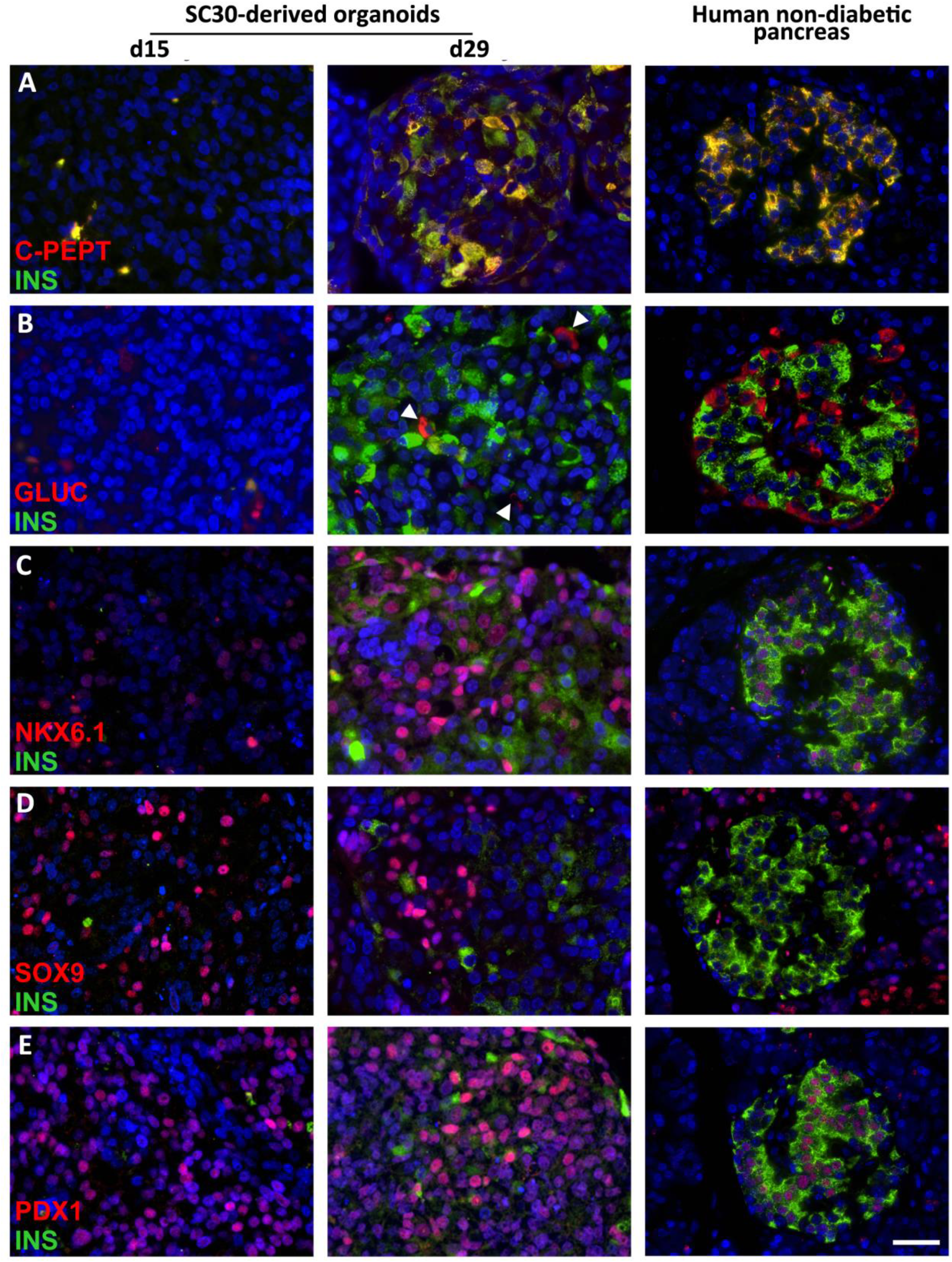
Immunhistochemical analysis of SC-derived pancreatic organoids. d15 spheroids and d29 stem cell-derived organoids derived in 3D from the SC30 clone were fixed, sectioned and double-stained for (**A**) C-peptide (red) and insulin (green) or insulin (green) and glucagon, NKX6.1, SOX9 or PDX1 (**B-E**, all in red). A human non-diabetic pancreas was taken as control. Scale bar = 20 µm.

**Figure 7.**
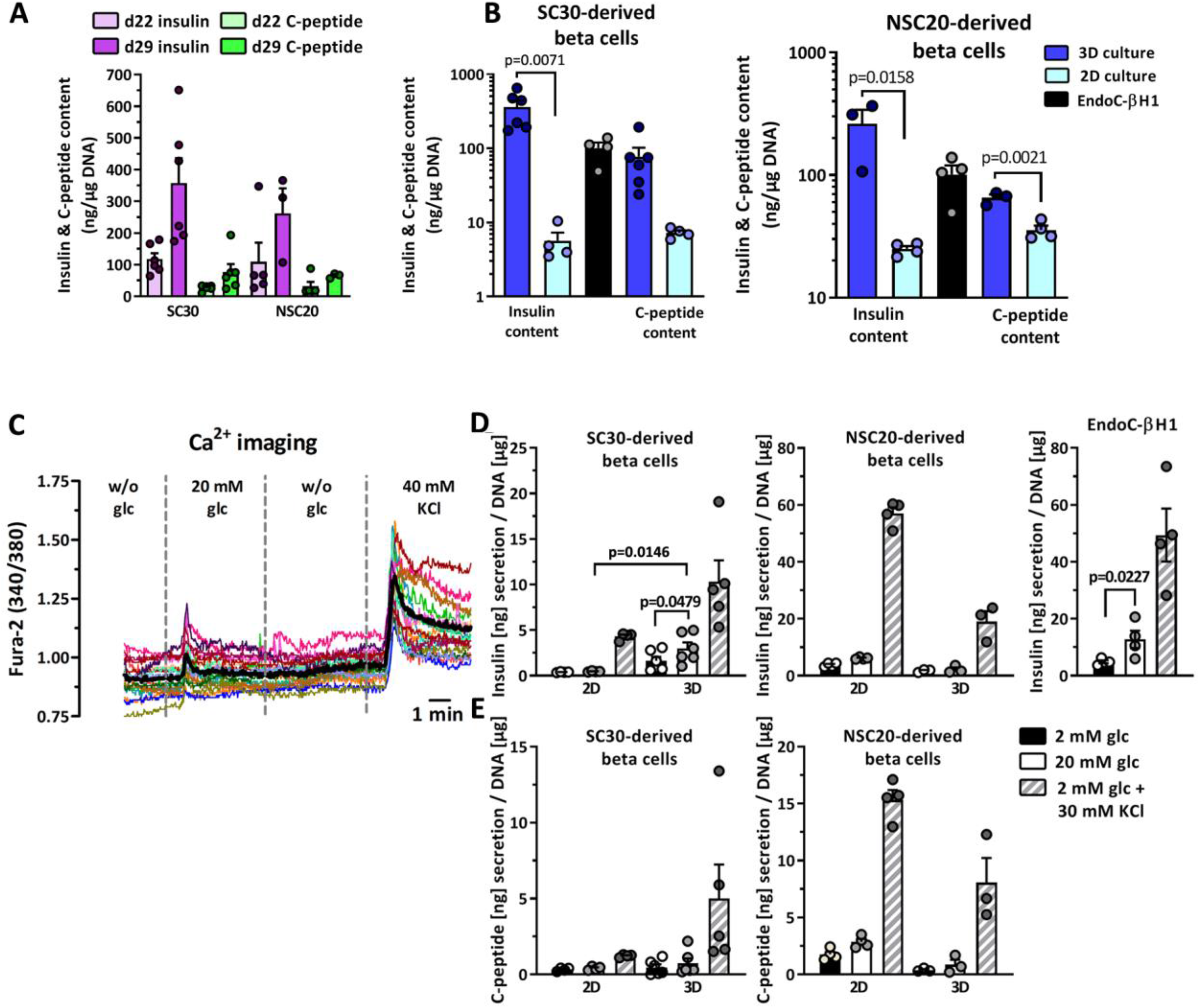
Insulin and C-peptide content. (**A**) Insulin and C-peptide content of NSC20- and SC30-derived organoids at d22 and d29 of differentiation in 3D. Values are means ± SEM, n= 3-6. (**B**) Insulin and C-peptide content of NSC20- and SC30-derived organoids at d29 differentiated in 2D or 3D in comparison to EndoC-βH1 cells. (**C**) Real time detection of cytosolic free-Ca^2+^ in SC30-derived organoids by recording of the Fura-2/AM emission ratio at 340 and 380 nm. The cells were perifused with basal KR w/o glucose, 20 mM glucose in KR, basal KR w/o glucose and finally KR plus 40 mM KCl. Mean value of 19 recorded cells are shown in bold black. (**D/E)** Measurement of insulin and C-peptide secretion in NSC20- and SC30-derived organoids at d29 after 2D and 3D differentiation in comparison to EndoC-βH1 cells. Data are means ± SEM, n= 3-6. Two-tailed *Student’s* t-test, ** p < 0.01, * p < 0.05.

Then the changes in free cytosolic Ca^2+^ were measured in real time by Fura-2AM assay. SC-derived organoids derived from SC30 and NSC20 were perifused with modified KR in the absence or presence of 20 mM glucose and 40 mM KCl. The SC30 cell clone showed a detectable increase in free calcium in the cytosol after exposure to glucose and potassium chloride (Fig. 7C), while the NSC20 cell clone only responded to potassium chloride (Supplementary Fig. 11).

Finally, the question was addressed whether 3D culture and differentiation could also improve glucose-induced insulin secretion (GSIS) in both cell lines (Fig. 7D/E). SC30-derived organoids showed a significant increase in insulin release when subjected to 20 mM glucose in a static assay. The insulin-releasing properties were also significantly improved compared to the 2D culture. NSC20-derived organoids showed a more robust insulin release in 2D culture though but neither in 2D or 3D the cells responded appropriately in response to glucose. EndoC-βH1 cells were measured as controls (Fig. 7E).

## Discussion

Here we report the generation and characterization of new hPSC reporter lines with insertion of the GFP2 and H-2K^k^ reporter genes following the *SOX9* open reading frame and an additional cell line with a knock-in of mCherry into the *INS* locus. Aided by these cell lines we established a new differentiation protocol geared towards SOX9 MPCs to efficiently drive hPSCs by 3D orbital shaking culture into SC-derived organoids with a beta cell content of > 40%. The SOX9 reporter cell lines showed peak SOX9/GFP2 expression after 10-13 days of differentiation using an experimental 2D differentiation protocol. This protocol is based on previous publications by our group in which we established a robust method for generating NKX6.1-/ PDX1+ pancreatic-duodenal cells [7, 8, 17] as well as adopted stage 3/stage 4 media described by Nostro and colleagues [11].

After purifying the GFP2+/GFP2-fractions, we were able to show that the GFP2+ cell population expressed high levels of *HNF6, NKX6-1, PDX*1 and *SOX9*, and thus are potentially SOX9 MPCs. Further analysis of MACS-purified SOX9 MPCs at d15 (end of stage 4) revealed mosaic-like expression of NKX6.1 and a few CPA1+ cells. Possibly these cells are representatives of the trunk region, which physiologically represents the niche for further endocrine development. Analysis of the MPC surface markers CD142, CD200 and GP2 revealed partial identity with MPCs generated with different protocols [12, 13].

By pursuing further down the developmental pathway and following the protocol published by Pagliuca and colleagues [14], the cells showed at d18 (end of stage 5) not only a more homogeneous NKX6.1 expression but also scattered NEUROG3+ cells. Then SOX9 MPCs readily developed into insulin/C-peptide-positive and glucagon-positive cells embedded in CK19+ epithelial cells. This fits in with findings from earlier studies in rodents that CK19+ fetal epithelium marks a source for endocrine islets [18]. In line with other studies polyhormonal cells were readily detected in 2D [19]. The SC-derived beta cell fraction generated in 2D showed no increased insulin secretion after glucose stimulation. A purification of SOX9 MPCs could not compensate for this deficit in function, although insulin and C-peptide content were increased, which confirms the effectiveness of an enrichment strategy [13].

Before the transition to 3D differentiation we systematically tested various compounds and conditions in stage 3 and 4 to increase the number of SOX9 MPCs. We can confirm that a 24 h pulse with stage 3 medium, which contains high FGF10 and a SHH inhibitor, is decisive for the differentiation into SOX9 MPCs. Withdrawal of nicotinamide and EGF in stage 4 also greatly reduced the number of SOX9 MPCs [11]. Interestingly, addition of the PKC activator PDBu was of no help at this stage of differentiation [20].

Regarding the transcriptional regulation, it has been reported, that SOX9 maintenance is controlled by Wnt/beta-catenin signaling [21], FGF-signaling via FGFR2b [22], Notch-signaling [23] and positive autoregulation [24]. Moreover, SOX9 and Wnt/beta-catenin form a regulatory loop and inhibit each other’s transcriptional activity. In chondrocytes SOX9 inhibits canonical Wnt-signaling by direct protein interaction with beta-catenin, yielding in inhibition and degradation of the protein [25, 26]. Vice versa Wnt/beta-catenin represses SOX9 gene expression in osteoblasts [27]. In our *in vitro* differentiation approach we can show for both hPSC lines that active canonical Wnt-signaling is an effective inhibitor of SOX9 MPC generation. This underlines the importance of this signaling pathway for differentiation protocols of hPSC into SC-derived beta cells. According to current and previous data, Wnt/beta-catenin not only prevents development of endocrine progenitors [28], but also earlier development of PDX1+ pancreatic-duodenal cells [8] and, as shown here, development into SOX9 MPCs.

Surprisingly our data indicate that differentiation into SOX9 MPCs is most effective in presence of EGF and not the commonly used growth factors FGF2/7 or FGF10 [13, 16, 29, 30]. FGF7 and FGF10 were less effective with regard to the absolute number of SOX9 MPCs or showed lower gene expression of typical MPC marker genes. Previously we had identified FGF2 as a repressor of early development into PDX1 pancreatic-duodenal cells [8]. In this study SOX9 MPC generation was also slightly less effective when compared to EGF. This was additionally evident from the reduced expression of *SOX9* and *NKX6-1* in FGF2-treated MPCs. In contrast to mouse studies for which an FGF10/FGFR2B/SOX9 feed-forward loop was described [22], the EGF-signaling pathway seems to play a greater role during differentiation into SOX9 MPCs in the human system.

3D differentiation in shaking orbital cultures or small bioreactors has become the standard for many somatic cell types [31-33]. Transition from 2D to 3D alone, without changes in extrinsic or other factors, can lead to a considerable phenotypic improvement [34]. At the same time, monitoring the *in vitro* differentiation with molecular biological methods such as RT-qPCR, immunofluorescence or Western blot is time-consuming, cumbersome and uneconomical. For these reasons, we converted the 2D differentiation protocol to 3D. In parallel, another knock-in was carried out into the *INS*-locus in order to be able to optimize the differentiation by means of flow cytometry and to easily follow the conversion of SOX9 MPCs into SC-derived beta cells. The 3D reconstructed islet-like organoids comprised SC-derived beta cells with important beta cell features such as expression of typical genes, insulin/C-peptide positive cells, very few polyhormonal cells, calcium influx after glucose exposure, and glucose-stimulated insulin secretion. Our results also show that transition from 2D to 3D culture not only results in a quantitative advantage, but also in an qualitatively improvement. The transition from 2D to 3D alone increased the insulin content in non-sorted cells by more than 50-fold (SC30). It is also important to note that we observed line-specific effects. While the hESC-based clone differentiated into glucose-responsive cells, this was not achieved for the iPSC-based clone. This result is probably attributed to cell line-specific barriers, which have already been described and represent a major obstacle in the establishment of patient-specific cell replacement therapies [35, 36].

The currently prevailing differentiation protocols were optimized towards the generation of mainly PDX1+/NKX6.1+ double-positive cells as the seed for SC-derived beta cells [14, 16]. Our protocol is based on an efficient generation of definitive endoderm using low activin A concentrations (> 90% endoderm cells), differentiation into PDX1+ pancreatic-duodenal cells by BMP and Wnt-inhition in presence of all-trans retinoic acid (∼80%) [7, 8], and optimized conditions to generate SOX9 MPCs (∼70%). After using the stage 5–7 media described by Pagliuca and co-workers [14], these cells effectively differentiated into SC-beta cells (> 40% positive cells). Thus, the new protocol that we present in this report may offer an alternative route to generate SC-derived beta cells.

Reporter cell lines are excellent tools to advance research into efficient differentiation methods [37-40]. The SOX9 reporter cell lines reported here are also an excellent model for studying the transition of human bipotent ductal/endocrine precursors into NEUROG3+ endocrine precursor cells. In view of the pleiotropic functions of SOX9 during development and tissue maintenance in diverse organs such as chondrocytes, testes, heart, lung, bile duct, retina and the central nervous system [25, 41, 42], these reporter cell lines are also suited for research on other matters. Additionally, the reporter gene knock-in into the *INS*-locus enables an exclusive look at insulin-producing cells and can therefore bypass the problem of a heterogenous composition of *in vitro* differentiated cells caused by not fully effective differentiation protocols.

## Materials & Methods

### Human cell culture

The human PSC lines Hes-3 (‘ESC’) and Phoenix (‘iPSC’, MHHi001-A) [43] were cultured on cell culture plastic coated with hESC-qualified Matrigel (Corning, Amsterdam, Netherlands). mTeSR1 (Stem Cell Technologies, Cologne, Germany) or StemMACS™ iPS-Brew XF medium (Miltenyi Biotec, Bergisch Gladbach, Germany) was used. Passaging was performed once a week in a ratio of 1:20 up to 1:40 and cluster were seeded onto fresh Matrigel-coated 6-well plates. EndoC-βH1 cells were cultured according to the standard protocol [44].

### Generation of hPSC reporter cell lines

To introduce DNA double-strand breaks (DSBs) into the genomic loci of *SOX9* and *INS* the CRISPR/Cas9 system was used. Putative sgRNAs were calculated with CCTop *(https://cctop.cos.uni-heidelberg.de/)* [45].The sgRNA for the *SOX9* locus was cloned into the pLKO5.U6 vector [46] and the sgRNAs, two nickase pairs, for the *INS* locus were cloned into the pX335-U6-Chimeric_BB-CBh-hSpCas9n vector [47] (Supplementary table 1). Reporter genes were introduced by HDR [48]. A scheme of the repair vectors is presented in Figure 1A/4A. Briefly 2 x 10^6^ hPSCs were nucleofected with 2 µg repair vector and 2 µg Cas9/sgRNA vector using the Neon Nucleofection System. Transfected hPSCs were seeded and selected after 24 h using either hygromycine b or blasticidin. Cell clones were picked after 10-14 days, expanded and genotyped by PCR and sequencing upon correct insertion of the transgenes into the desired loci.

### 2D experimental differentiation protocol

For differentiation of hPSCs in 2D, hPSC colonies were dissociated into single cells by Trypsin/EDTA (T/E) (Biochrom, Berlin, Germany) and centrifuged for 3 min at 300 x g. The pellet was re-suspended in mTeSR1 or iPS-Brew XF containing 5 µM Y-27632 (Selleck Chemicals, Munich, Germany) and a defined number of cells (100,000 cells/12-well plate cavity, 250,000 cells/6-well plate cavity and 1.3-1.45×10^6^ cells per 10 cm cell culture dish) were seeded on Matrigel-coated cell culture plastics. Cells were allowed to re-attach and differentiation was initiated after 24 h. Differentiation was performed according to an adopted 7-stage protocol (Figure 1B/2A) [7-9, 11, 14]. Media compositions were as followed: stage 1a medium (24 h), RPMI1640, (Biochrom) plus 0.5fold B27 supplement (Thermo Fisher Scientific, Schwerte, Germany), 1% penicillin/streptomycin (P/S) (penicillin: Santa Cruz Biotechnology, Dallas, USA; streptomycin: Sigma-Aldrich, Munich, Germany), 2 mM glutamine, 1-fold non-essential amino acids (NEAA, Thermo Fisher Scientific, Schwerte, Germany), 1 mM sodium pyruvate (Capricorn scientific, Ebsdorfergrund, Germany), 0.5-fold ITS-X (Thermo Fisher Scientific) 0.25 mM vitamin C, 30 ng/ml activin A (Stem Cell Technologies), and 3 µM CHIR99021 (Cayman Chemical, Ann Arbor, USA); stage 1b medium (72 h) was composed as stage 1a medium but lacked CHIR99021. The required activin A concentration was titrated for every individual lot (Supplementary Fig. 1A). Stage 2 medium (96 h), advanced RPMI1640 (Thermo Fisher Scientific) plus 0.5 fold B27 supplement, 1% P/S, 2 mM glutamine, 1x NEAA, 0.25 mM vitamin C, 5 ng/ml FGF7 (Stem Cell Technologies), 2 µM IWR-1 (Selleck Chemicals, Munich, Germany), 0.5 µM LDN193189 (Sigma-Aldrich) and 1 µM all-trans retinoic acid (ATRA, Sigma-Aldrich); stage 3 medium (24 h), DMEM (Biochrom) plus 1% P/S, 2 mM glutamine, 0.5 fold B27 supplement, 50 g/ml vitamin C, 50 ng/ml FGF10 (Stem Cell Technologies), 0.25 µM Sant-1 (Selleck Chemicals), 2 µM ATRA, 0.5 µM LDN193189; stage 4 medium (6 days), DMEM plus 1% P/S, 2 mM glutamine, 0.5 fold B27 supplement, 50 g/ml vitamin C, 10 mM nicotinamide (Sigma-Aldrich), 200 ng/ml EGF (Stem Cell Technologies), 0.5 µM LDN193189; stage 5 medium (3 days), BE5 stock medium plus ITS-X (1:200), 10 µg/ml heparin (Sigma-Aldrich), 20 ng/ml betacellulin (Stem Cell Technologies), 10 µM ZnSO_4_(MerckMillipore, Schwalbach, Germany), 0.25 µM Sant-1, 50 nM ATRA, 1 µM XXI (Stem Cell Technologies) or 1 µM LY411575 (Selleck Chemicals), 10 µM Alk5iII (Santa Cruz Biotechnology, Dallas, USA) or 10 µM RepSox (Selleck Chemicals), 1 µM GC1 (Tocris, Bristol, United Kingdom), 3 nM staurosporine (Cayman Chemical), 5 µM Y-27632, 100 nM LDN193189; stage 6 medium (4 days) was composed as stage 5 medium but lacked ATRA and Sant-1; stage 7 medium (7 days), CMRLM stock medium plus 20 nM insulin (Sigma-Aldrich), 10 µM ZnSO_4_, 10 µg/ml heparin, 15 µM ethanolamine (Sigma-Aldrich), 10 µM Trolox (Sigma-Aldrich), 10 µM Alk5iII or RepSox, 1 µM GC1, 70 nM apo-transferrin (Sigma-Aldrich), medium trace elements A (1:1000, Corning), medium trace elements B (1:1000, Corning), chemically defined lipid concentrate (1:2000, Thermo Fisher Scientific). BE5 stock was composed of MCDB131 (Thermo Fisher Scientific) plus 20 mM D-(+)-Glucose, 1.754 g/l NaHCO_3_, 2% FAF-BSA (SERVA, Heidelberg, Germany), 2 mM glutamine and 1% P/S. CMRLM stock was composed of 2% FAF-BSA, 1% P/S, 2 mM glutamine and 5 mM sodium-pyruvate. FGF2, FGF7, FGF10 (each 100 ng/ml) and IWP4 (1 µM) were obtained from StemCell Technologies. The PKC activator PDBu was purchased from Tocris (biotechne, Minneapolis, USA). Unless otherwise mentioned, chemicals were obtained from Riedel-de Haen, (Munich, Germany), J.T. Baker (Chihuahua, Mexico) or Sigma-Aldrich.

### 3D production protocol

Differentiation of hPSCs in 3D was initiated by seeding hPSCs on Matrigel-coated cell culture plastics. Media compositions remained the same as described for 2D with slight differences. Stage 3 and stage 4 medium were supplemented with 2 µM IWR-1 and 100 ng/ml EGF was supplemented to stage 4 medium. Differentiation proceeded in 2D according to the 7-stage protocol (Figure 1B/2A) until day 12 of differentiation. Then the cells were washed with PBS, dissociated into single cells by T/E and centrifuged for 4 min at 300 x g. The cell pellet was re-suspended in 0.1 mg/ml DNaseI grade II (Sigma-Aldrich) in PBS + 10% FCS and incubated at room temperature for 15-20 min. Then the cells were centrifuged for 4 min at 300x g and re-suspended in stage 4 medium supplemented with 5 µM Y-27632. 1.5-2.5 x 10^6^ cells/ml were seeded on a 6-well suspension culture plate (Greiner bio-one, Kremsmünster, Austria) and cultivated at 100 rpm and 25 mm hub on an orbital shaker (Infors HT, Celltron, Bottmingen, Switzerland) according to the 7-stage protocol. The medium was changed daily until day 12 of differentiation and thereafter every second day.

### Western Blot

Cells at d0, d4, d8, d15, d18 and d29 were taken up in PBS and sonified. A protease inhibitor mixture (Roche Diagnostic, Mannheim, Germany) was then added. The protein content was determined by BCA assay (Thermo Fisher Scientific). 40 µg of total protein was loaded and separated by SDS-PAGE and transferred by electro-blotting to a PVDF membrane. Blocking was performed with 5 % nonfat dry milk in PBS plus 0.1 % Tween 20. The membrane was incubated with anti-CPA1 (1:1000, Origene, cat# TA500053, clone OTI2A3) overnight at 4°C then washed and followed by incubation with the peroxidase-labeled secondary antibody for 1 h. As a loading control actin was used. Protein bands were visualized by chemiluminescence using the detection kit (GE Healthcare Europe, Solingen, Germany) on a chemiluminescence imager (INTAS Science imaging, Göttingen, Germany).

### Gene expression analysis

Isolation of total RNA was carried out using the Machery&Nagel Nucleospin RNA plus Kit (Macherey&Nagel, Düren, Germany). cDNA was synthesized from 500-2000 ng total RNA using RevertAid™ H Minus M-MuLV Reverse Transcriptase (Thermo Fisher Scientific) and random hexamer primers. cDNA samples were then diluted to 2.5-5 ng/µl and measured in a qPCR reaction with the GoTaq^®^ qPCR Master Mix (Promega, Walldorf, Germany). All reactions were performed by a 2-step PCR in triplicates followed by melting curve analysis on a ViiA7 real-time PCR cycler (Thermo Fisher Scientific). Primers are specified in Supplementary table 2. Data normalization was performed with qBasePlus (Biogazelle, Zwijnaarde, Belgium) against the geometric mean of the housekeeping genes *G6PD, TBP* and *TUBA1A*. RT-qPCR data are presented as calibrated normalized relative quantities (CNRQ). Analysis of housekeeping gene stability was performed with the geNorm algorithm.

### Flow cytometry

Cells were washed with PBS and dissociated using T/E. Organoids from 3D culture were collected in a 15 ml conical tube, centrifuged at 50 x g for 5 min and subsequently dissociated by incubation with gentle cell dissociation solution (StemCell Technologies) for 15 min and additional T/E for 10 min. Single cells were then centrifuged at 300 x g for 3 min and re-suspended in PBS + 2% FCS before flow cytometric measurement. For flow cytometric staining 1 x 10^6^ cells were washed, incubated for 20 min at 4°C with primary conjugated antibodies and washed twice prior to analysis. Flow cytometric measurements were performed on a CyFlow ML flow cytometer (Partec, Münster, Germany). Data analysis was performed using the FlowJo software (Ashland, OR, USA).

The following conjugated antibodies were used: anti CXCR4-PE (FC15004, Neuromics, Minneapolis, USA), anti CXCR4-APC (130-098-357, Miltenyi Biotec), anti-H-2K^K^-APC (130-117-324, Miltenyi Biotec), Anti-CD177-APC (130-101-512, Miltenyi Biotec), Anti-CD275-APC (130-098-738, Miltenyi Biotec), anti-CD200-APC (130-118-203, Miltenyi Biotec), anti-CD142-PE-Vio616 (130-115-720, Miltenyi Biotec). Anti-GP2 (D277-3, MBL international, Woburn, MA, USA) was stained at 1:500 and then labelled with 1:500 diluted anti-mouse AF647 (Dianova, Hamburg, Germany).

### Cell sorting

Fluorescence activated cell sorting (FACS) was performed at the central facility of the Hannover Medical School. For MACS 1 x 10^7^ dissociated cells were taken up in PBE buffer (PBS, pH 7.2, 0.5 % BSA, and 2 mM EDTA) and were then conjugated with anti H2-K^K^ magnetic microbeads (Miltenyi-Biotec) for 15 min on ice. Cell sorting was then performed on an autoMACS Pro (Miltenyi-Biotec).

### Immunofluorescence

For immunofluorescence staining, hPSCs were seeded onto Matrigel-coated glass cover slides (SPL Life Sciences, Pocheon, South Korea) with 5 µM Y-27632. After 24 h the cells were fixated with 4 % (w/v) paraformaldehyde (PFA), buffered in PBS, pH 7.4. The same fixation was used for organoids at day 15 and day 29 after differentiation which were embedded in paraffin and sectioned. After pretreatment and blocking steps the same primary antibodies were used for cells and organoids and incubated for 1-3 h at room temperature or overnight at 4°C (Supplementary Table 3). The cells as well as organoids were stained with conjugated either with AlexaFluor or Cy fluorophores secondary antibodies (Dianova) and counterstained with mounting medium containing DAPI. For comparison immunostaining of human islets from four non-diabetic donors were performed (for details see new (Supplementary Table 4). Stained cells or stained organoids were examined using an inverse Olympus IX81 microscope (Olympus, Hamburg, Germany) or an upright Olympus microscope BX61 and representative pictures were taken of each analyzed sample as previously described [49].

### Insulin and C-peptide content and secretion

Cells grown in 2D or 3D culture were washed with bicarbonate-buffered Krebs-Ringer (KR) solution and hungered for 2 h in KR without glucose, supplemented with 0.1 % albumin. Thereafter the cells were stimulated either with 2 or 20 mM glucose or 2 mM glucose and 30 mM KCl for 1 h. To measure insulin and C-peptide secretion, the medium was removed and centrifuged for 5 min at 700 x g. For this measurement, the cells were taken up in PBS, sonicated and centrifuged for 5 min at 700 x g. The resulting supernatant was used to determine hormone secretion. Secreted insulin in the supernatant and insulin content of the incubated cells were determined by radioimmunoassay using human insulin as standard and the resulting values were normalized to DNA content [44]. Human C-peptide content and C-peptide secretion was measured by a sandwich ELISA assay (DRG Diagnostics, Marburg, Germany).

### Calcium imaging

Cytosolic free-Ca^2+^ was determined with Fura-2/AM. Day 28 clusters were dissociated, grown on Matrigel-coated glass coverslips overnight and loaded with 3 μM Fura-2/AM by incubation in modified Krebs-Ringer (KR) solution (25 mM HEPES, 3 mM glucose, and 1.5 % BSA) for 30 min at 37°C. Perifusion was performed with a modified KR solution containing 0 mM or 20 mM glucose and 0 mM glucose plus 40 mM KCl at a flow rate of 1 ml/min using a peristaltic pump (Ismatec, Zürich, Switzerland). Images were taken every 2 sec using the inverted IX81 microscope equipped with an UPlanSApo 40×0.95 numerical aperture objective (Olympus) and an incubation chamber to maintain 60% humidity, 37°C, and 5% CO_2_, with excitation/emission filter settings of 340±26 nm & 387±11 nm, respectively.

### Statistics

Unless stated otherwise values represent mean ± SEM and the number of independent experiments (n= independent biological replicates) is stated in each figure legend. Statistical analyses were performed using the GraphPad Prism analysis software (Graphpad, San Diego, CA, USA) using unpaired, two-tailed *Student’s* t-test or ANOVA plus *Dunnett’s* or *Tukey’s* post-hoc tests for multiple comparisons. P-values for *Student’s* t-test are depicted in each figure. A summary of all GraphPad Prism statistical test results in particular the ANOVA plus post-hoc tests are available online.

## Acknowledgements

This work has been supported by the Deutsche Forschungsgemeinschaft (DFG, German Research Foundation, NA 1285/2-1). The MHHi001-A cell line was kindly provided by Dr. A. Haase from the LEBAO (Leibniz Research Laboratories for Biotechnology and Artificial Organs), Hannover Medical School. We gratefully acknowledge the technical assistance of Rebecca Chucholl and Monika Funck. We would also like to acknowledge the assistance of the Cell Sorting Core Facility of the Hannover Medical School supported by the Braukmann-Wittenberg-Herz-Stiftung and the DFG.

## Supplementary data

**Supplementary figure 1.**
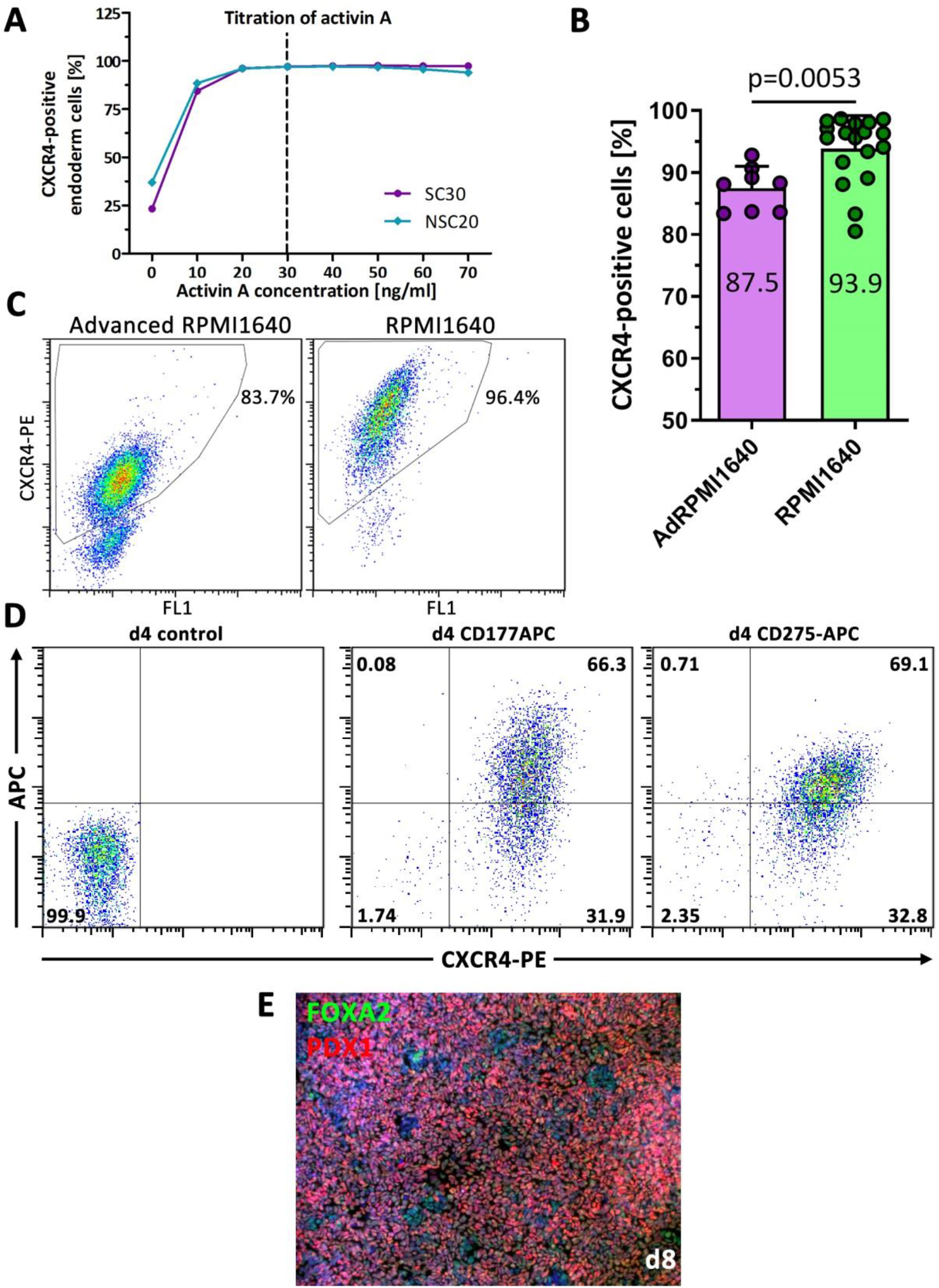
Endoderm and pancreatic-duodenal differentiation efficiency. (**A**) Titration of the optimal activin A concentration for differentiation of hPSC into CXCR4-positive endoderm cells. (**B**) Flow cytometric quantification of CXCR4 at d4 of endoderm differentiation. Data are means ± SEM, n= 8-18, two-tailed *Student’s* t-test, ** p < 0.01. Differentiation protocol based on [9]. (**C**) Representative flow cytometry dot plots of CXCR4 staining in two endoderm differentiation media. (**D**) Double flow cytometric staining of CD177-APC/CXCR4-PE and CD275-APC/CXCR4-PE at d4 of endoderm differentiation. (**E**) PDX1/FOXA2 double-positive pancreatic duodenal cells at d8 of differentiation. Differentiation protocol based on [7] and [8].

**Supplementary figure 2.**
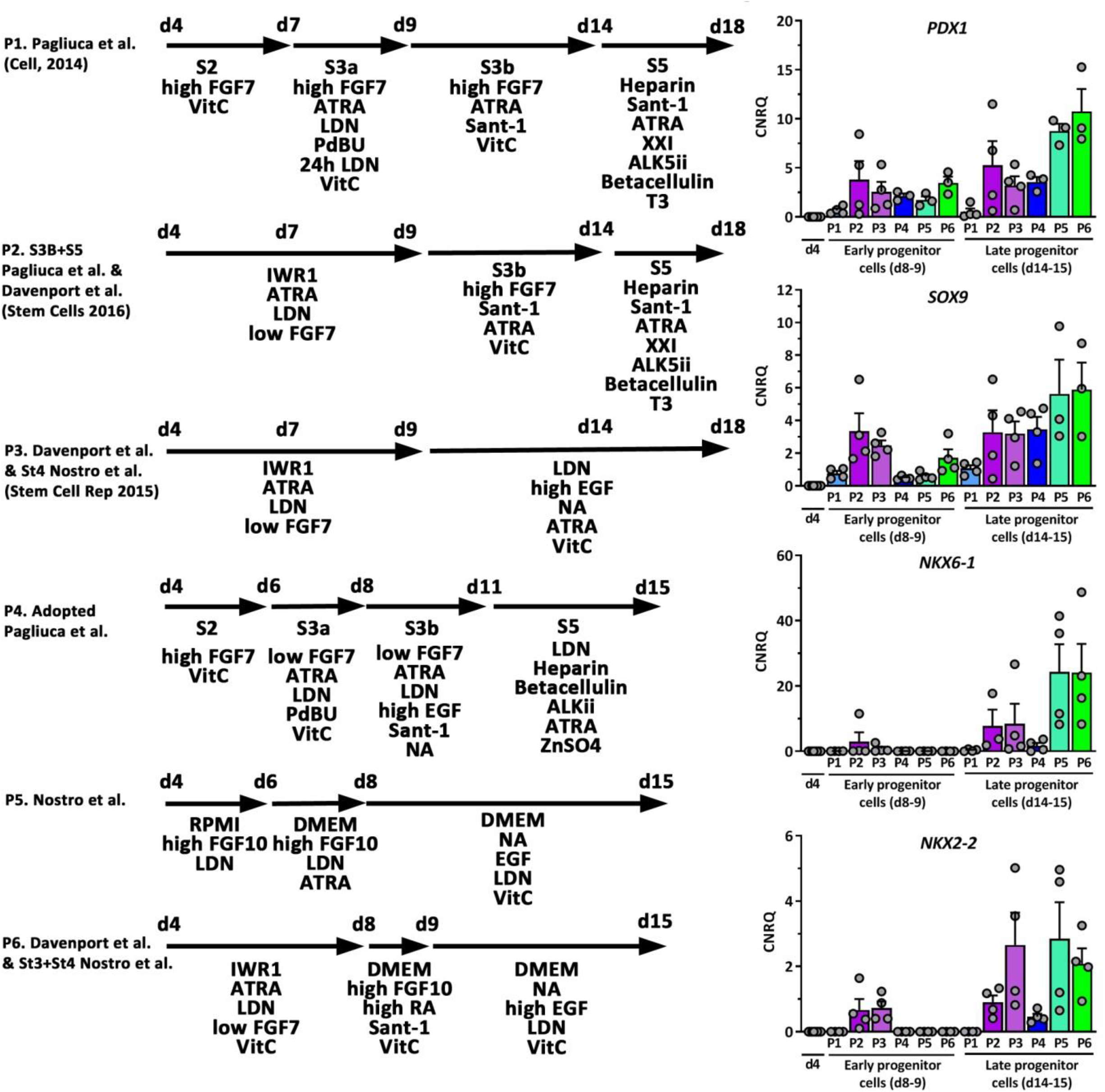
Comparison of six adopted differentiation protocols for the generation of MPCs from hPSC. Depicted is the relative gene expression of *PDX1, SOX9, NKX6-1*, and *NKX2-2*. Data are means ± SEM. n= 3-4. The differentiation protocols were adopted from [7], [8], [11] and [14].

**Supplementary figure 3.**
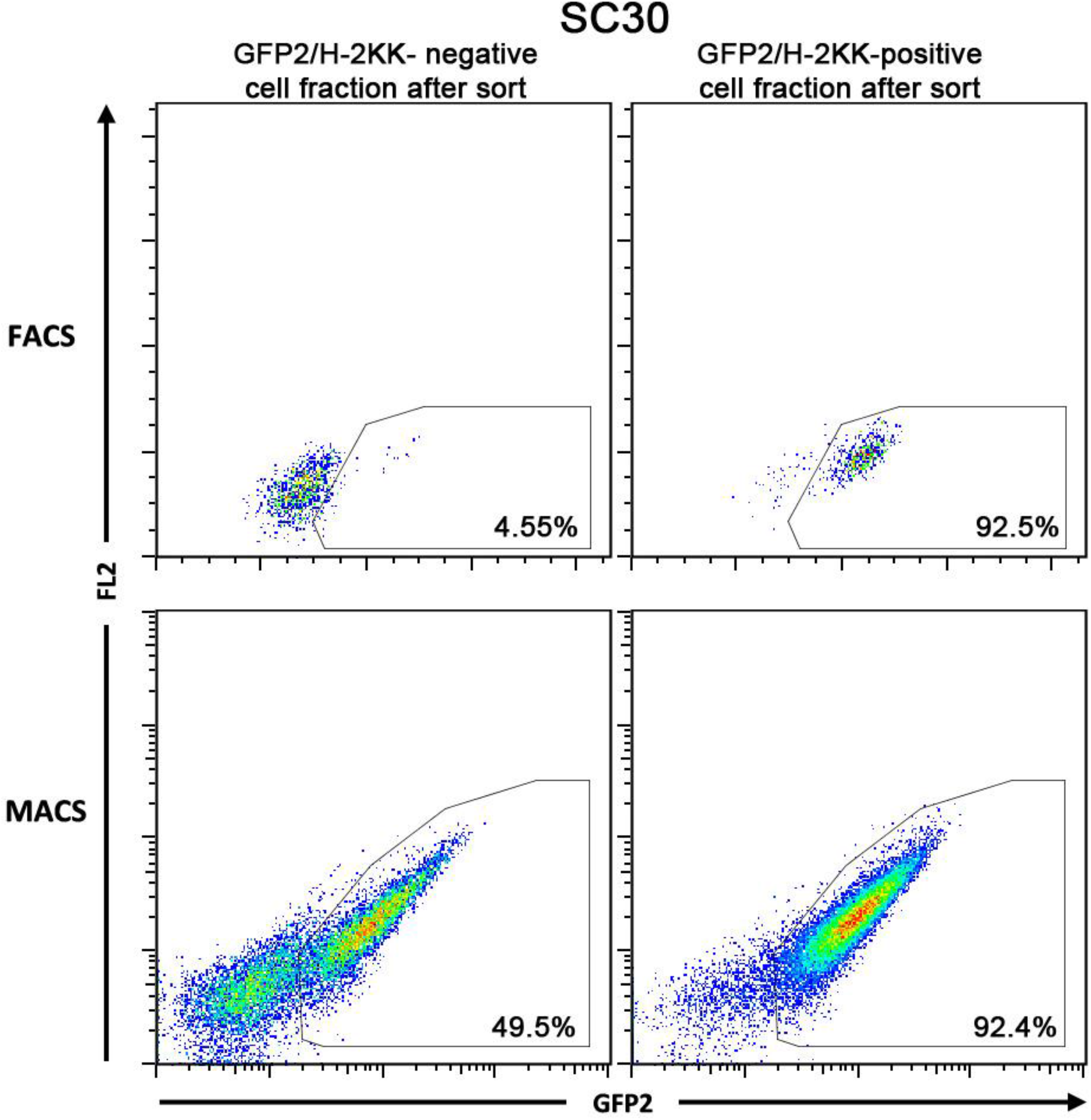
Representative dot plot presentation of cell sorting experiments using the SC30 cell clone. GFP2^+^ pancreatic progenitors can be sorted by FACS (upper images) or MACS (upper images) with comparable efficiencies.

**Supplementary figure 4.**
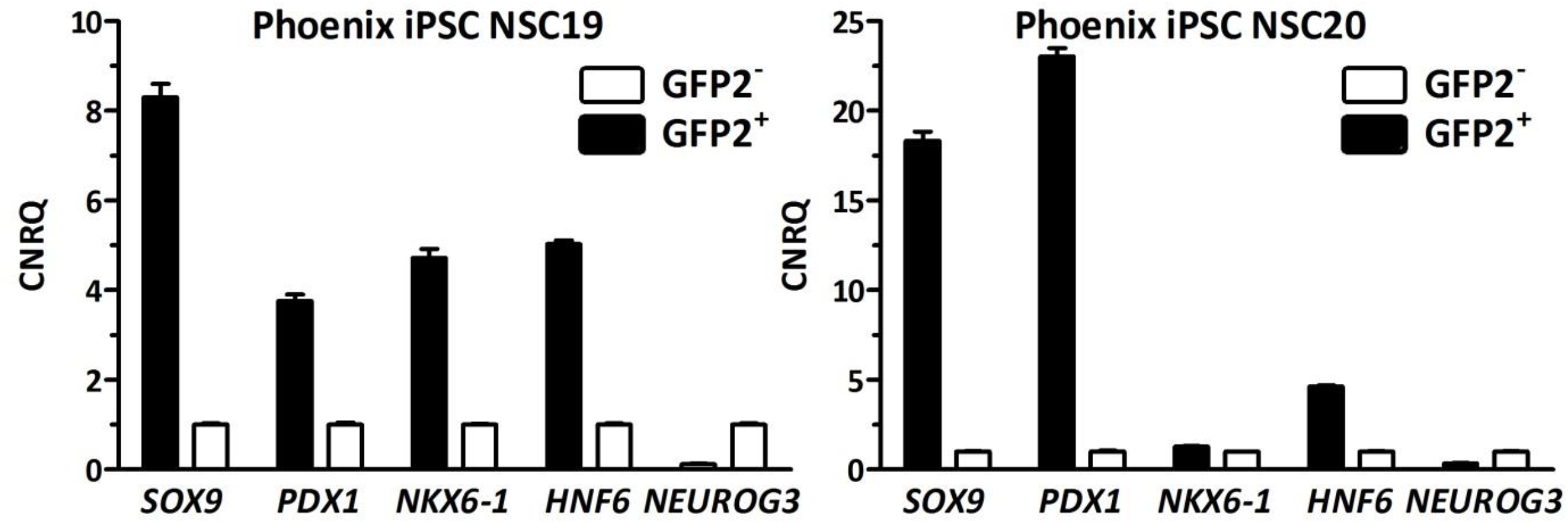
RT-qPCR analysis of sorted NSC19 and NSC20 derived GFP2+ and GFP2-cells at d12 of differentiation. Depicted is the relative gene expression of *SOX9, PDX1, NKX6-1, HNF6*, and *NEUROG3*. Data are means ± SD from a single sorting experiment measured in triplicate.

**Supplementary figure 5.**
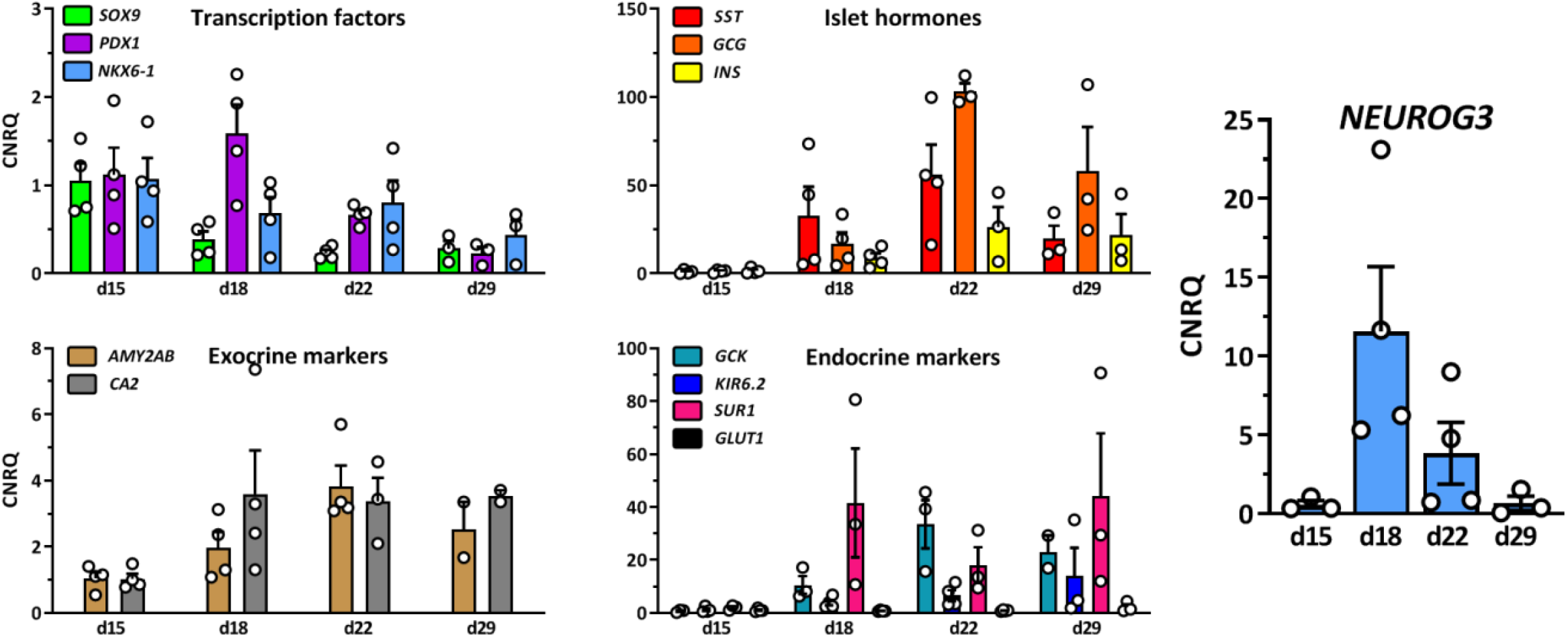
RT-qPCR analysis of sorted GFP2^+^ cells after MACS at day 12 of differentiation using the SC30 cell clone. Further differentiation was conducted according to the 2D experimental protocol. Depicted is the relative gene expression of the transcription factors, *SOX9, PDX1, NKX6-1* and *NEUROG3*, the islet hormones somatostatin (*SST*), glucagon (*GCG*) and insulin (*INS*), the exocrine marker genes amylase 2 (*AMY2AB*) and carbonic anhydrase 2 (*CA2*) and the endocrine marker genes glucokinase (*GCK*), *KIR6*.*2, SUR1* and *GLUT2*. Data are means ± SEM, n= 2-4. Data are normalized to housekeeping genes and d15 samples scaled to 1.

**Supplementary figure 6.**
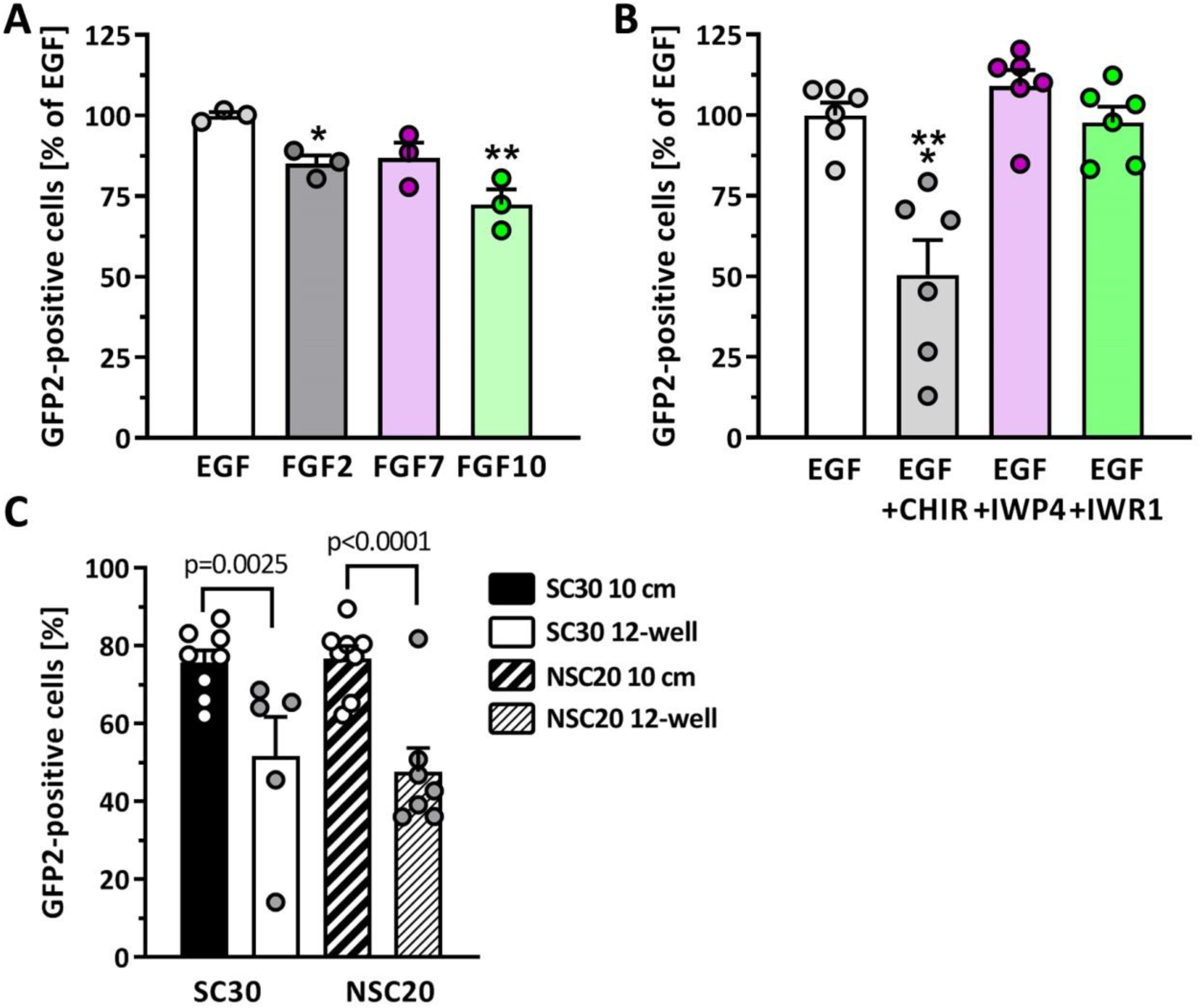
Effect of growth factors, Wnt/beta-catenin signaling and upscaling on the generation of GFP2+ pancreatic progenitors. (**A**) Effect of different growth factors each used at 100 ng/ml on GFP2 expression in NSC20 cells. Data are means ± SEM. n= 3, two-tailed *Student’s* t-test, ** p < 0.01, * p < 0.05. (**B**) Effect of canonical Wnt-signaling on GFP2 expression in NSC20 cells. The pathway was activated by CHIR (3 µM) or inhibited by IWP4 (1 µM) or IWR-1 (2 µM). Data are means ± SEM, n= 6, two-tailed *Student’s* t-test, *** p < 0.001 (**C**) Flow cytometric quantification of GFP+ pancreatic progenitors from the cell lines SC30 and NSC20 at d12 of differentiation differentiated in 12-well plate cavities or 10 cm cell culture dishes. Data are means ± SEM, n= 8-11. Two-tailed *Student’s* t-test, *** p < 0.001, ** p < 0.01.

**Supplementary figure 7.**
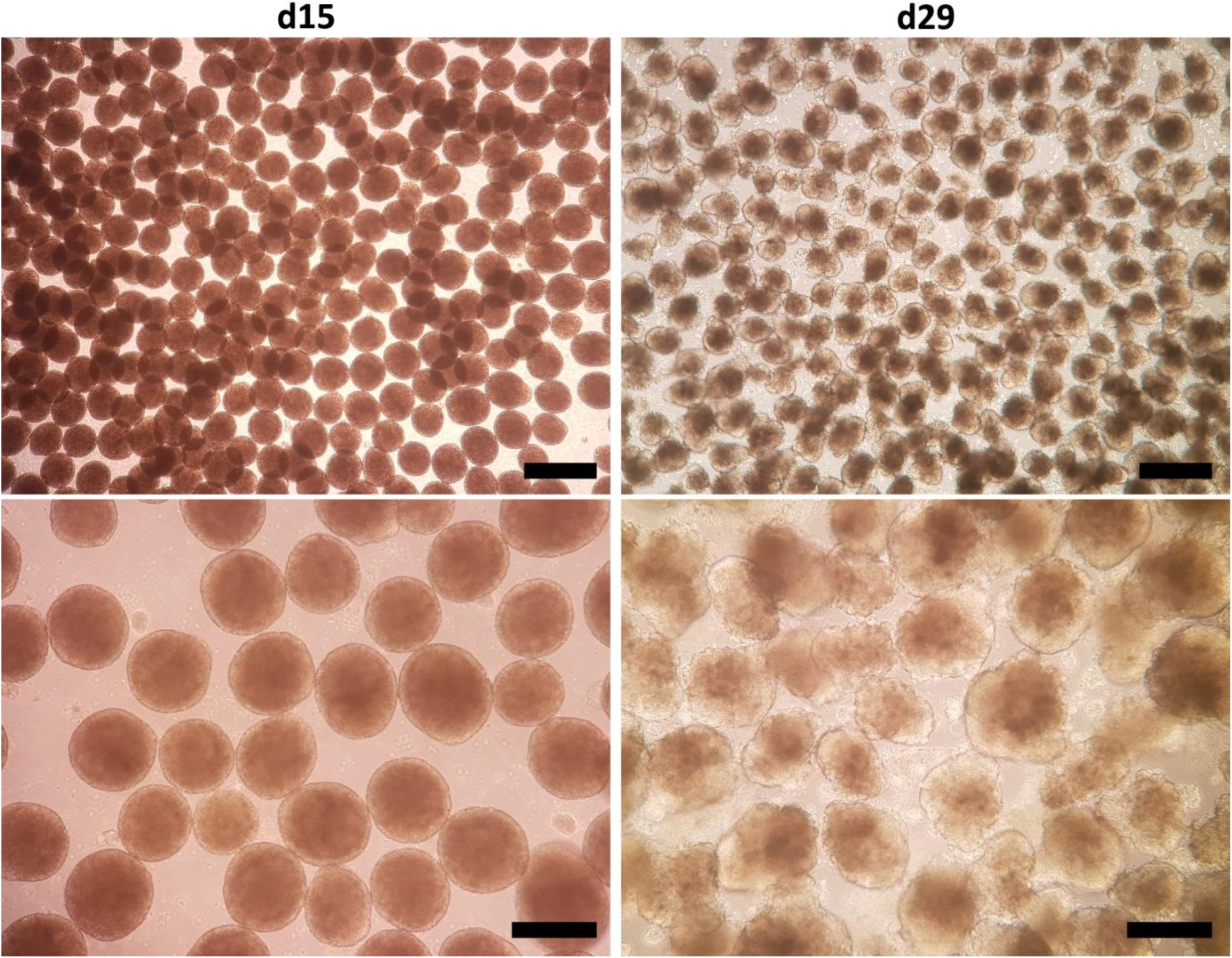
Generation of pancreatic spheroids (left images) and stem cell-derived organoids (right images) by 3D shaking culture. Shown are representative images of cell spheroids 3 days (left) and organoids 17 days (right) after transfer from 2D adherent culture to 3D orbital suspension culture at 100 rpm. Scale bar for lower magnification image = 500 µm. Scale bar for higher magnification image = 200 µm.

**Supplementary figure 8.**
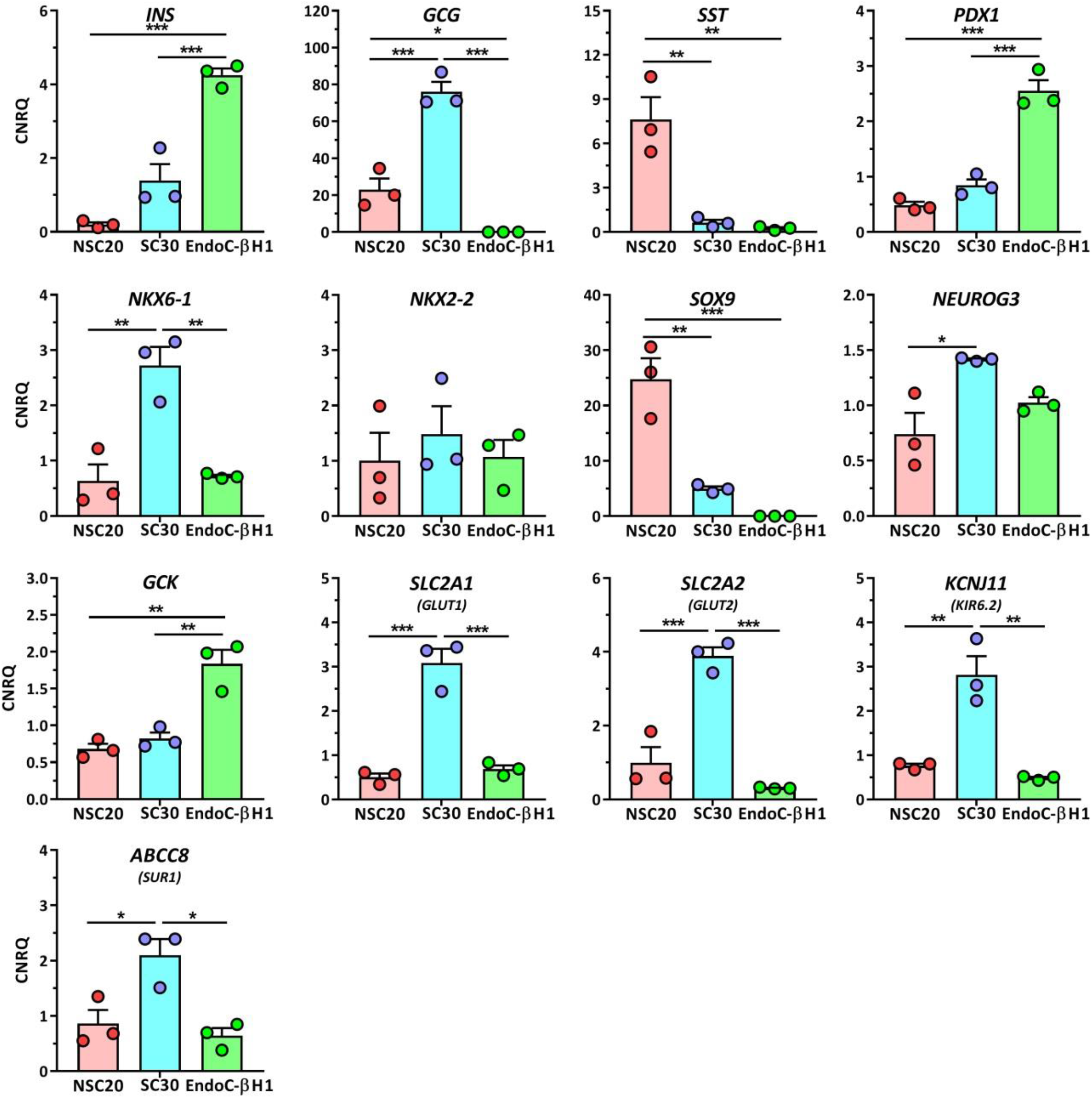
Relative gene expression of pancreatic and endocrine genes in NSC20-and SC30-derived organoids after 3D differentiation compared to EndoC-βH1 cells. Data are means ± SEM, n= 3. ANOVA plus *Tukey’s* post test, *** p < 0.001, ** p < 0.01, * p < 0.05.

**Supplementary figure 9.**
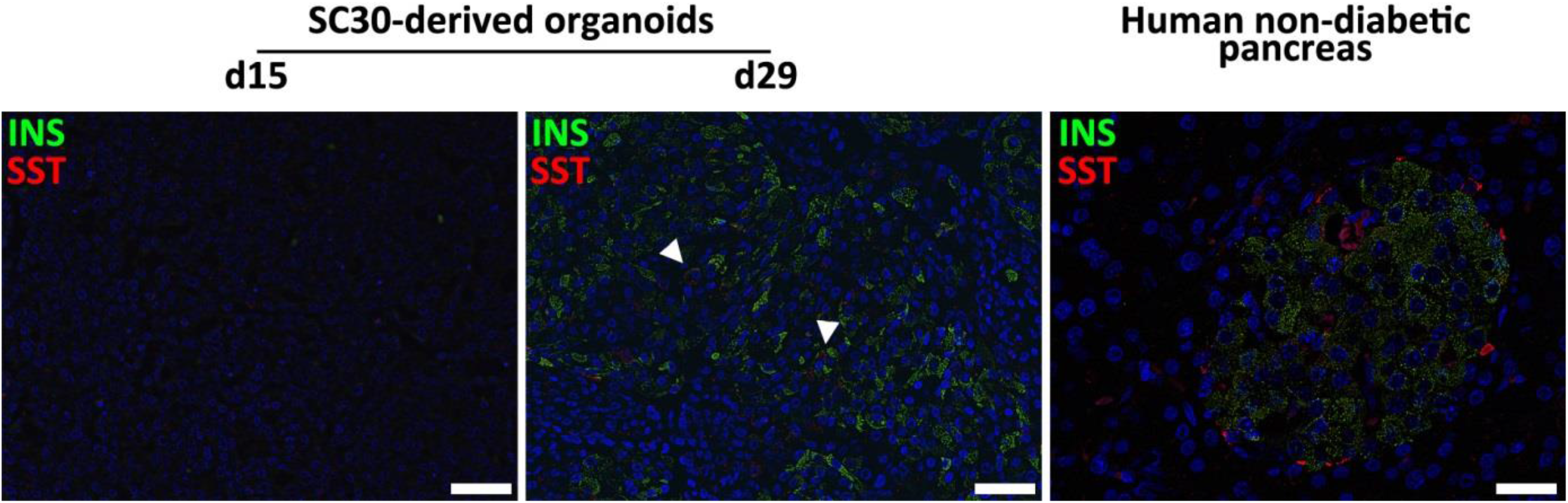
Immunhistochemical analysis of SC-derived pancreatic organoids. D15 spheroids and d29 stem cell-derived organoids derived in 3D from the SC30 clone were fixed, sectioned and double-stained for somatostatin (red) and insulin (green). A human non-diabetic pancreas was taken as control. Arrowheads mark polyhormonal cells. Scale Bar = 50 µm.

**Supplementary figure 10.**
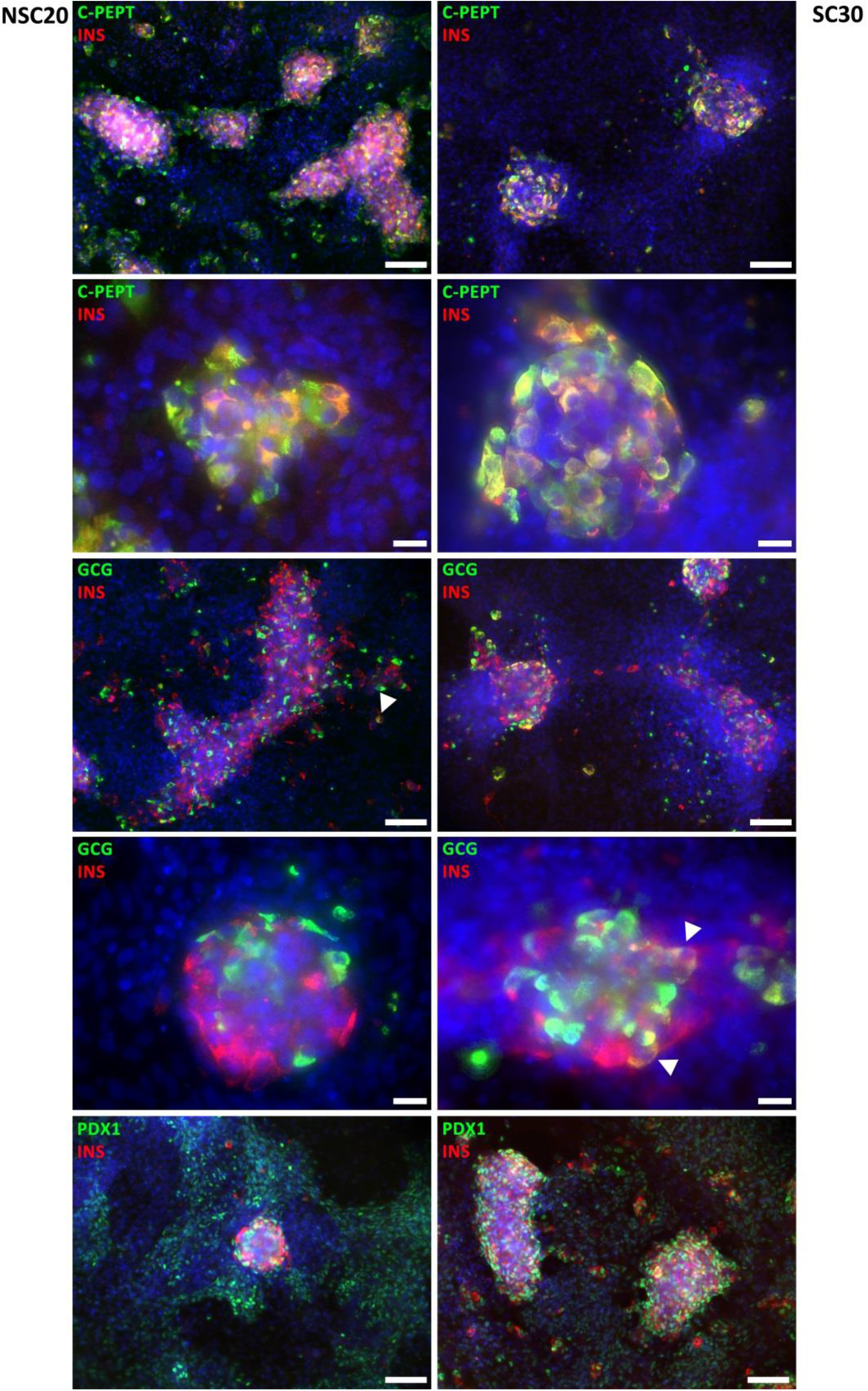
Immunofluorescence staining of NSC20-derived (left images) and SC30-derived (right images) pancreatic and endocrine cells at d29 using the production protocol in 2D. Double staining of insulin (red) and C-peptide, glucagon and PDX1 (all in green). Counterstaining with DAPI. Scale bar for lower magnification image = 100 µM, scale bar for higher magnification images = 20 µM. Arrowheads indicate polyhormonal insulin and glucagon co-expressing cells.

**Supplementary figure 11.**
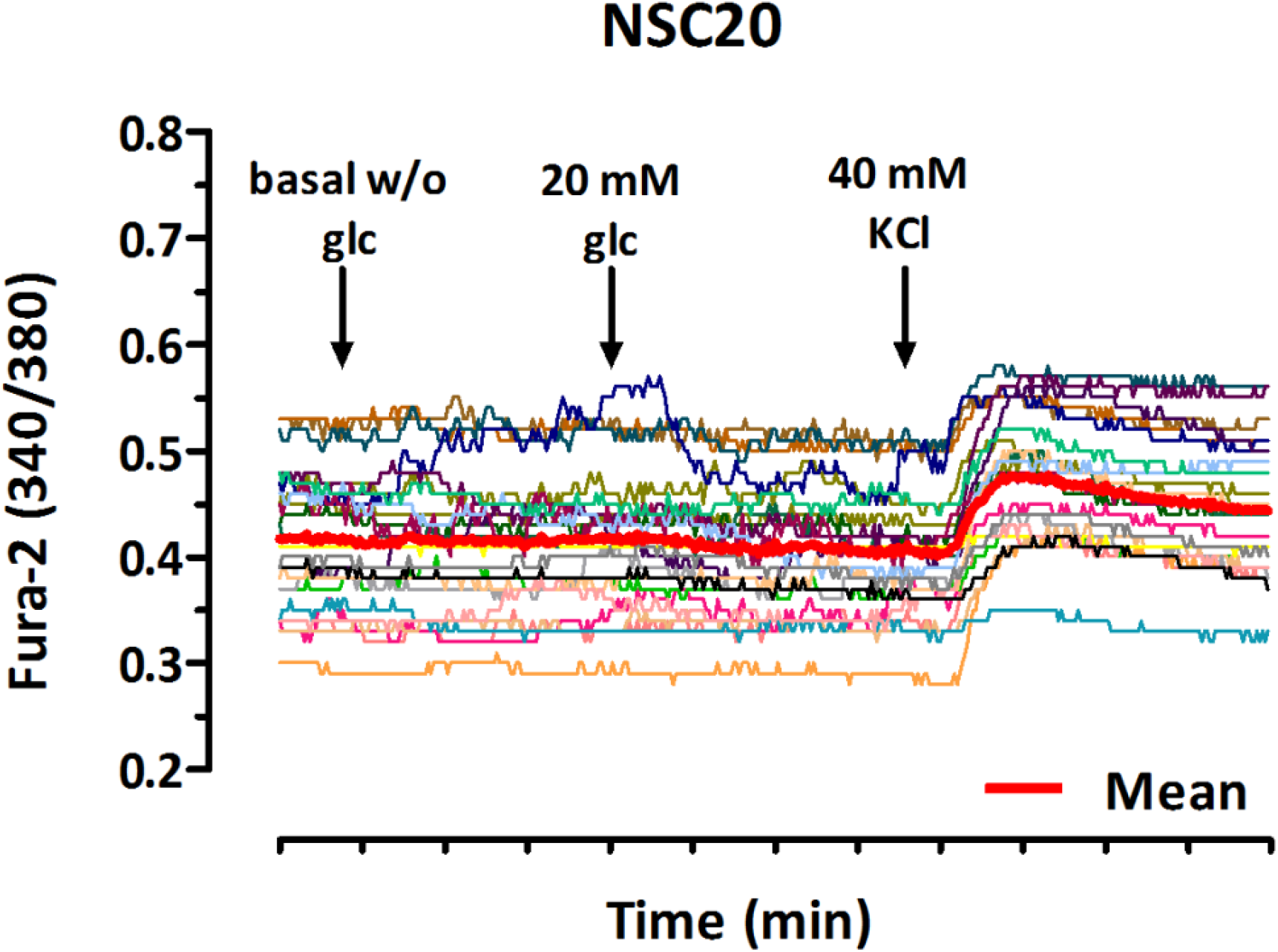
Recording of the Fura-2/AM emission ratio at 340 and 380 nm over 13 minutes. NSC20-derived organoids were dissociated, seeded on glass slides for 24 h and loaded with Fura-2/AM. Then the cells were stimulated with basal KR Ø glucose, 20 mM glucose in KR, basal KR Ø glucose and finally KR plus 40 mM KCl. Mean value of all 15 cells in bold red.

**Supplementary Table 1:**
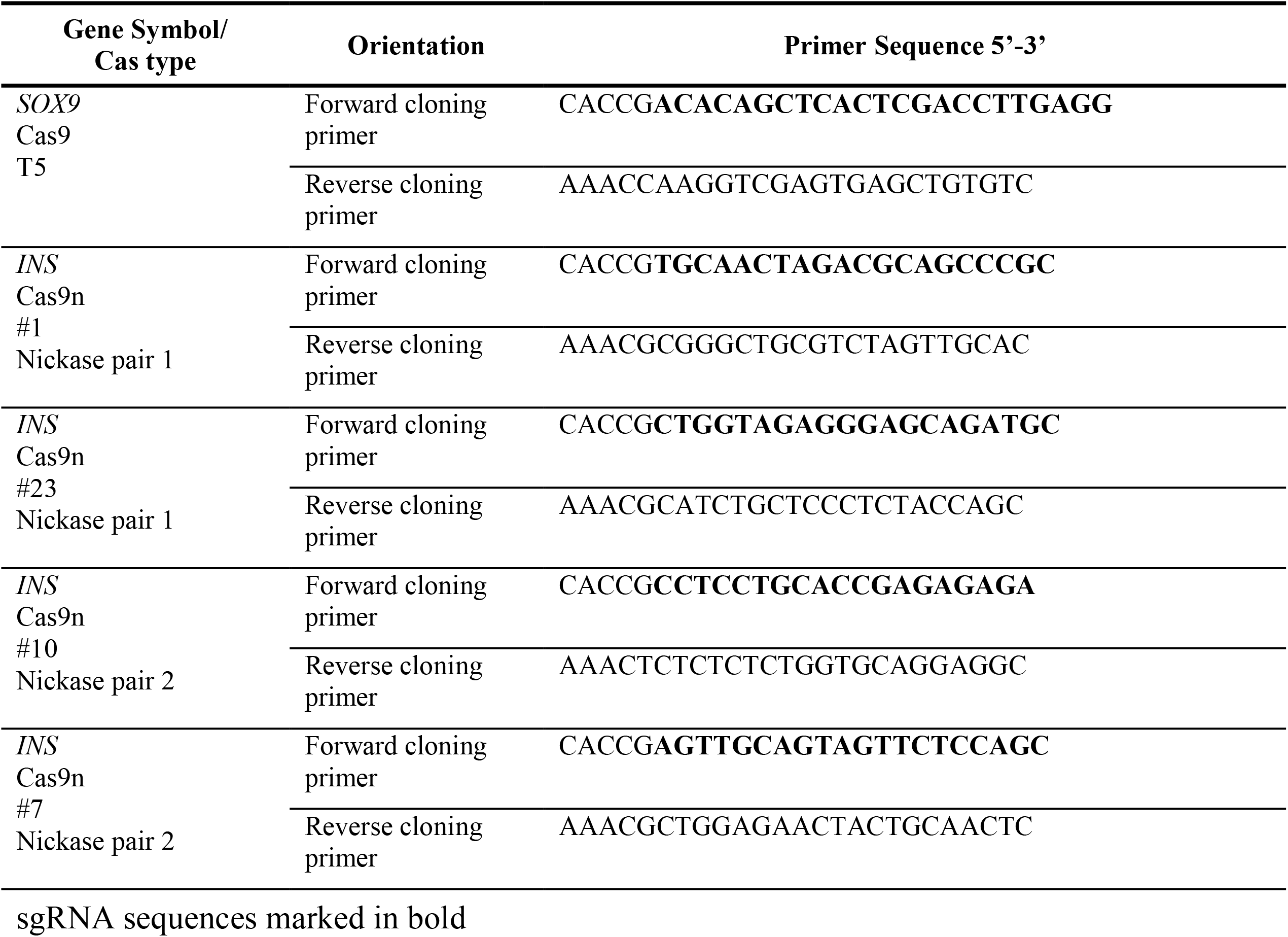
sgRNAs for HDR

**Supplementary Table 2:**
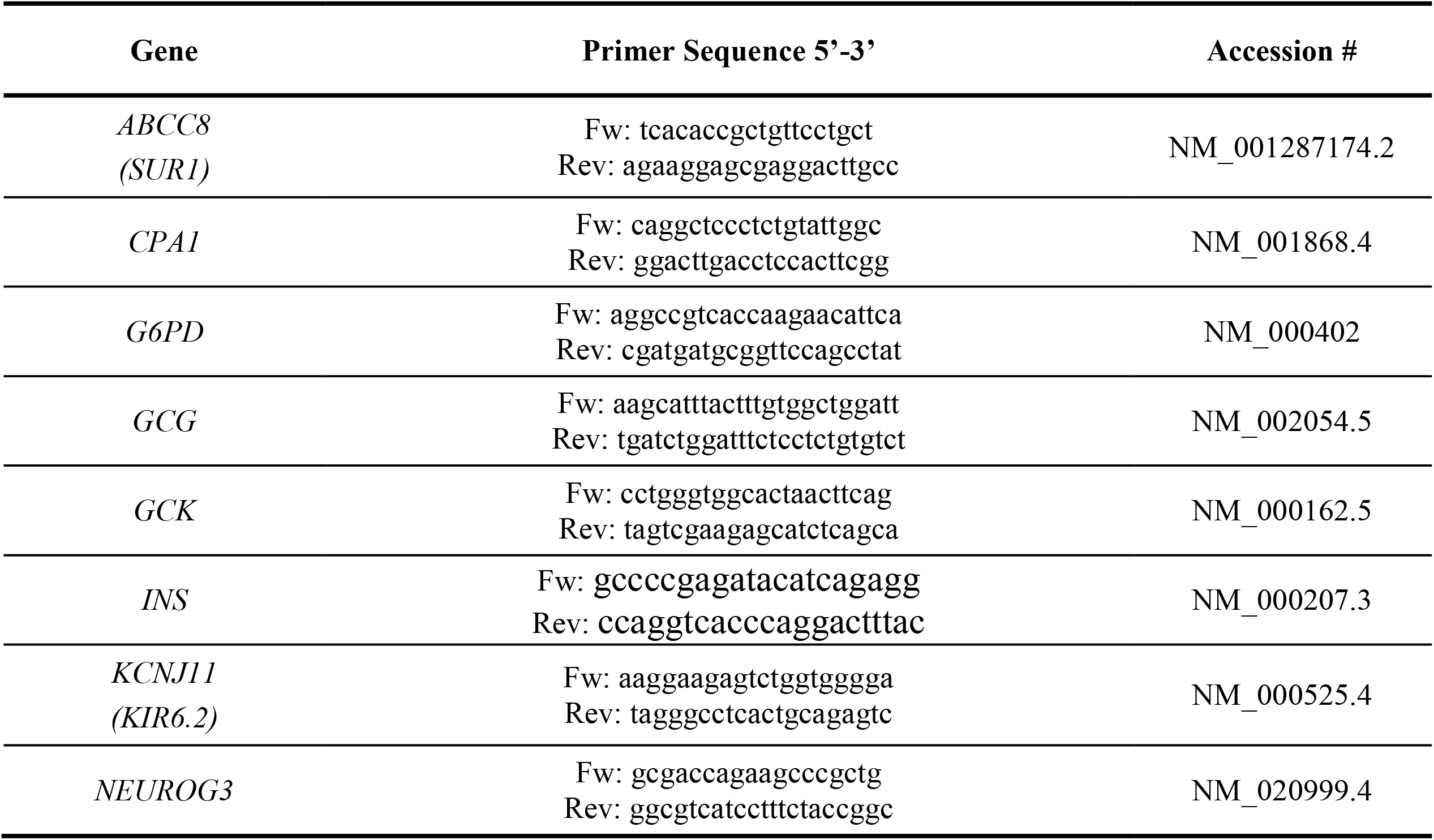
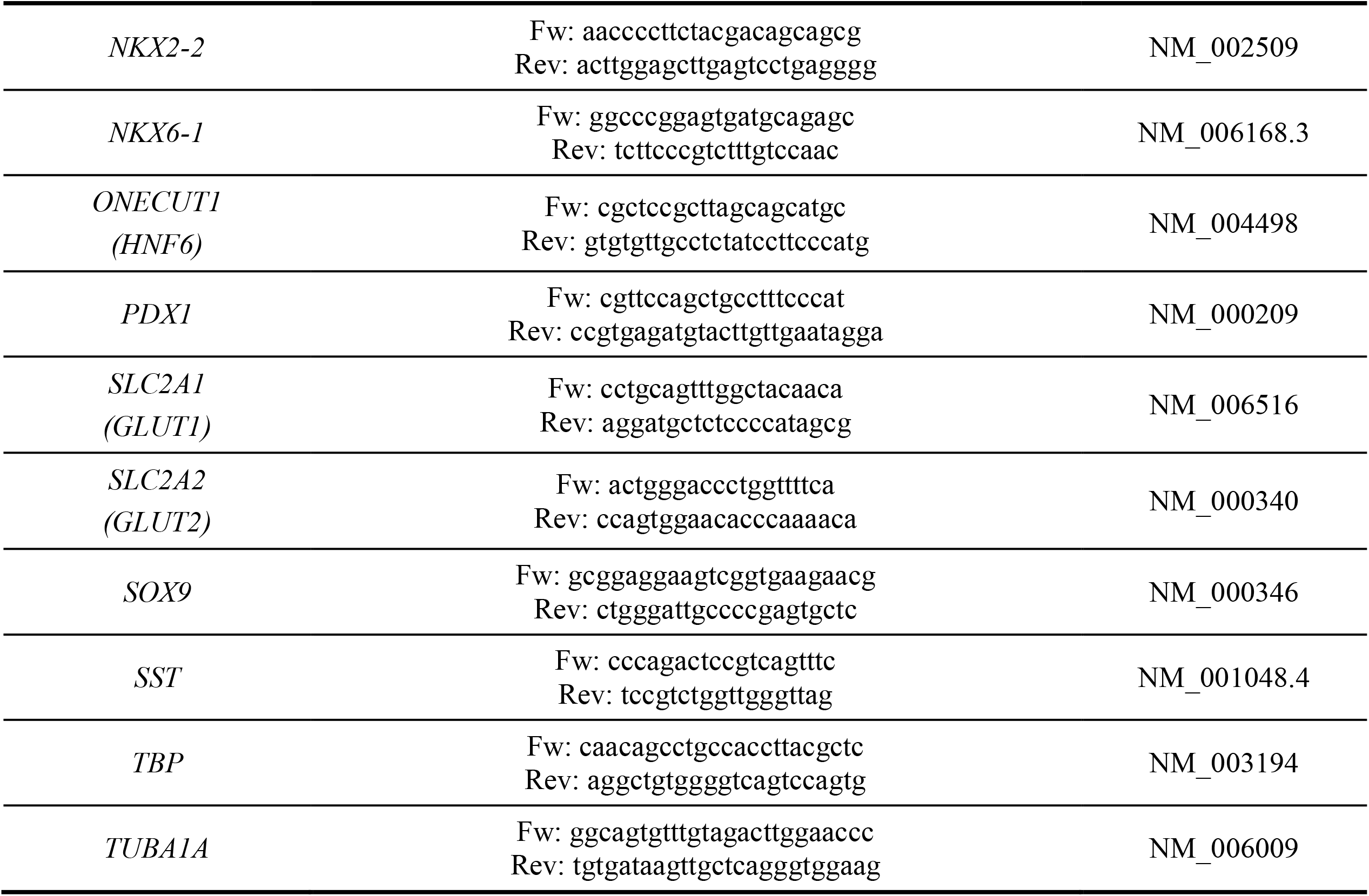
Primer pairs for gene expression analysis.

**Supplementary Table 3:**
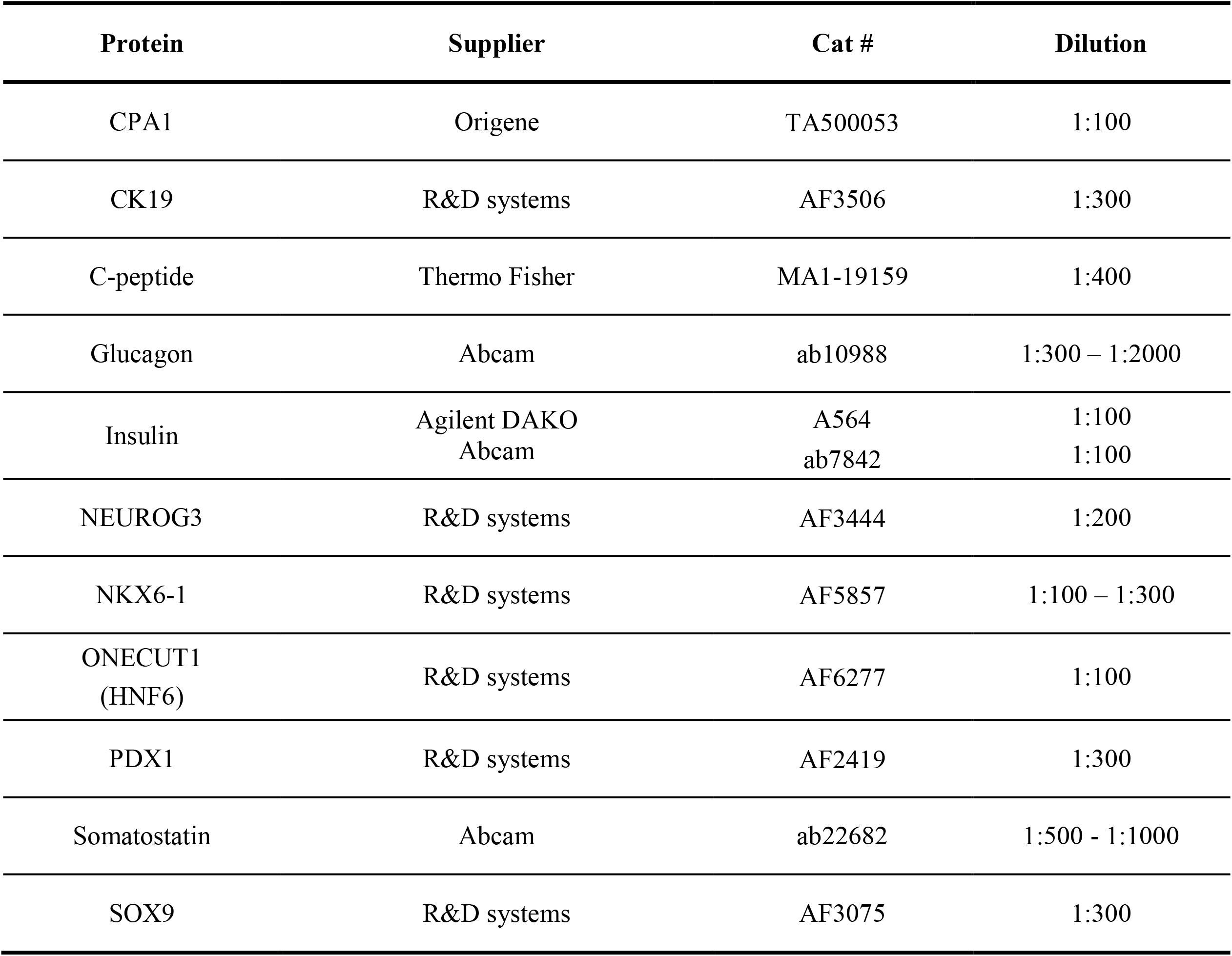
Antibodies used in this study.

**Supplementary Table 4:**
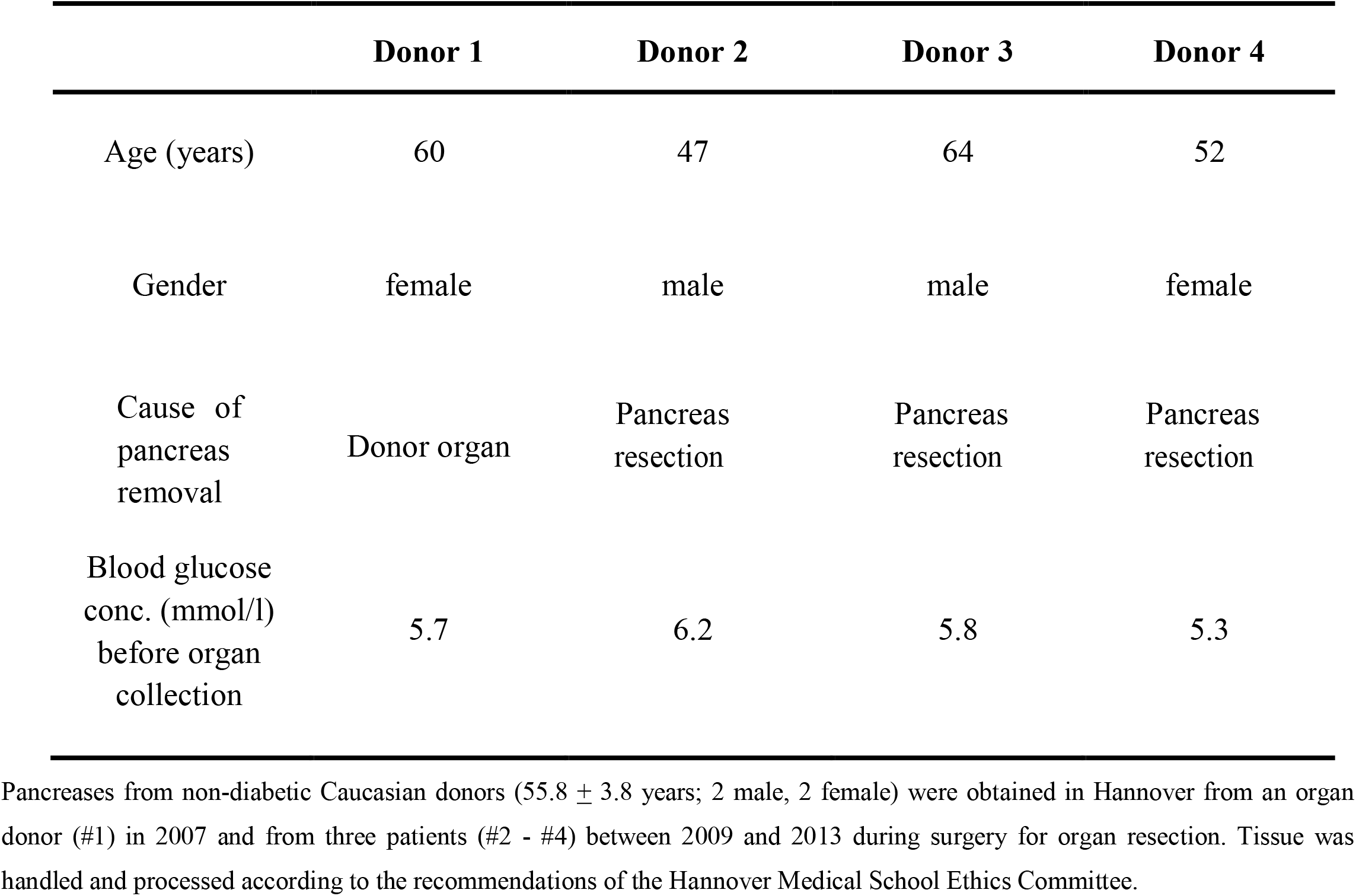
Data on the non-diabetic human pancreas organ donors

